# Urbanisation drives biodiversity loss, fitness decline and community shifts in cavity-nesting Hymenoptera

**DOI:** 10.1101/2025.07.28.667168

**Authors:** Atilla Çelikgil, Lars Henzschel, Jonathan Sänger, Pascal Dennis Krause, Adrian Nemetschek, Martin Husemann, Panagiotis Theodorou

## Abstract

Urbanisation is reshaping landscapes worldwide, posing challenges to biodiversity and ecosystem function. Cavity-nesting Hymenoptera, including bees and wasps, deliver vital ecological services but are increasingly vulnerable to urbanisation. In this citizen science-based study, we deployed 286 trap nests across Leipzig and Hamburg (Germany) to investigate how urbanisation and landscape-scale environmental features influence the diversity, community composition, reproductive success, offspring survival and demographic status of these insects. Using Bayesian hierarchical and phylogenetically informed modelling, we found that urbanisation significantly reduced species richness, reproductive output and offspring survival in both guilds, despite stable nesting activity, suggesting potential ecological trap effects. For bees, urbanisation also reduced the probability of nests functioning as demographic sources. Environmental drivers shaped responses in taxon-specific ways: bee richness and survival increased with landscape diversity but declined with fragmentation, while wasp richness and nesting declined with higher temperatures, though survival increased with more green cover. Bee nests were more likely to serve as demographic sources in higher temperatures. Community composition responses also diverged; bee communities shifted via increased species turnover, while wasp assemblages remained relatively stable. Temperature was a major driver of compositional dissimilarity in both groups, whereas landscape diversity was associated with homogenisation in bees and diversification in wasps. Our findings demonstrate that cities filter cavity-nesting Hymenoptera through multiple ecological pathways, highlighting the need for differentiated conservation strategies that consider both habitat structure and taxon-specific responses.

## 1. Introduction

Urbanisation is a dominant force reshaping ecosystems globally, transforming natural and semi-natural habitats into densely built environments through the expansion of infrastructure and human settlements (Grimm et al., 2008; Kundu & Pandey, 2020; Seto et al., 2012). As more than 60% of the global population is projected to live in urban areas by 2050 (Zhou et al., 2019), the ecological footprint of cities continues to grow, driving widespread land-use change and biodiversity loss (Huang et al., 2019). Urbanisation not only leads to habitat loss, fragmentation and degradation, but also fundamentally alters abiotic conditions, increasing impervious surface cover, pollution and urban heat island effects (McDonnell & Hahs, 2009). Together, these changes introduce a suite of complex and interacting environmental stressors that can disrupt ecological dynamics and influence evolutionary processes (Johnson & Munshi-South, 2017; McKinney, 2006; Seto et al., 2012; Theodorou, 2022).

Hymenoptera, a highly diverse order of insects that includes bees, wasps, sawflies and ants, are among the groups strongly affected by urban environmental change (Fenoglio et al., 2020; Liang et al., 2023). These insects play vital ecological roles as pollinators, biological control agents and decomposers, supporting plant reproduction, regulating insect populations and facilitating nutrient cycling across both natural and human-modified environments (Wilson, 1987). While numerous studies have examined urban effects on Hymenopteran species richness and abundance (Liang et al., 2023), the ecological consequences of urbanisation are highly context-dependent and the relative influence of key drivers (e.g., temperature, habitat fragmentation, resource availability (Baldock et al., 2019; Biella et al., 2022; Geppert et al., 2023; Theodorou et al., 2017; Theodorou, Herbst, et al., 2020; Theodorou, Radzevičiūtė, et al., 2020) remains unresolved. Importantly, most existing research has been limited in geographic scope, methodological consistency, or analytical depth, complicating our understanding of urbanisation’s generalisable effects on insect biodiversity (Banaszak-Cibicka & Żmihorski, 2012; Fenoglio et al., 2020; Theodorou, 2022).

Beyond taxonomic metrics, urbanisation also affects the composition of ecological communities, altering which species persist, in what combinations and with what consequences. These community composition shifts often reflect differential species responses to urban stressors, such as variation in habitat preference, thermal tolerance and resource specialisation (Aronson et al., 2016; McKinney, 2008). Urban communities often exhibit biotic homogenisation, favouring generalist, disturbance-tolerant taxa at the expense of habitat specialists (Liang et al., 2023; McKinney, 2006). Such shifts in community structure can mask functional losses even where species richness is unchanged. Moreover, shifts in community composition can have cascading effects on ecosystem functions such as pollination and biological control, ultimately influencing ecosystem resilience and the provision of services in urban landscapes (Alberti, 2015; Theodorou, 2022).

While changes in species community structure are increasingly recognised (Ayers & Rehan, 2023; Herrmann et al., 2023; Wilson & Jamieson, 2019), the fitness consequences of urbanisation remain largely underexplored. Yet, determining whether urban environments support not just species presence, but also successful reproduction, offspring survival and demographically viable nests is critical for assessing long-term population viability (Rivkin et al., 2019). Fitness outcomes, such as reproductive output, offspring survival and demographic status, offer mechanistic insights into whether cities act as ecological traps or as adaptive environments that can sustain biodiversity (Rivkin et al., 2019). Despite their importance, few urban ecological studies directly link landscape characteristics to reproductive fitness and demographic status (but see Dürrbaum et al., 2022; Moretti et al., 2021; Prendergast, 2023; Sexton et al., 2021; Theodorou et al., 2022), particularly across spatially replicated urban contexts, leaving a key gap in our understanding of how urban landscapes shape not only community structure, but also population-level processes.

Cavity-nesting Hymenoptera, including many solitary bees and wasps, offer a powerful model system to fill this gap (Staab et al., 2018; Tscharntke et al., 1998). Their reliance on discrete nesting substrates (e.g., hollow stems, deadwood, or artificial nests) makes them highly sensitive to habitat change and ideal bioindicators of urban environmental quality (Everaars et al., 2011; Staab et al., 2018; Tscharntke et al., 1998). Standardised trap nests (aka "insect hotels") allow for consistent monitoring across space and time, enabling detailed assessments of nesting behaviour, community composition, diet, parasitism, and crucially, reproductive success (Staab et al., 2018). Although numerous studies have used trap nests to examine urban impacts (Casanelles-Abella et al., 2022, 2024; da Rocha-Filho et al., 2020, 2022; Dürrbaum et al., 2022; Everaars et al., 2011; Fernandes et al., 2022; Fortel et al., 2016; Makinson et al., 2017; Moretti et al., 2021; Pereira-Peixoto et al., 2014, 2016; Prendergast, 2023; Sexton et al., 2021; Turo & Gardiner, 2021; Xie et al., 2022) findings have been inconsistent, particularly regarding species richness, offspring survival and reproductive output. For instance, the relationship between species richness and urban green space availability has ranged from negative (Makinson et al., 2017) to positive (Casanelles-Abella et al., 2024). Studies on reproductive output and offspring survival have also produced conflicting results, with some reporting increases in urban areas (Moretti et al., 2021; Sexton et al., 2021) and others reporting declines (Casanelles-Abella et al., 2024; Dürrbaum et al., 2022). These discrepancies likely stem from methodological variation, confounding environmental factors and limited replication across cities. To resolve these uncertainties, there is a pressing need for integrative, standardised, multi-city studies that assess multiple facets of biodiversity, including fitness, while accounting for key environmental drivers. Here, we address this gap by deploying 286 trap nests across diverse urban and peri-urban landscapes in two German cities, Leipzig and Hamburg, as part of a large-scale citizen science initiative. By integrating detailed landscape metrics with Bayesian hierarchical and phylogenetically informed modelling, our study provides a comprehensive, spatially explicit assessment of how urbanisation influences cavity-nesting Hymenoptera. Specifically, we ask: (1) Does urbanisation affect species richness, number of nests, reproductive output, offspring survival and demographic status in cavity-nesting bees and wasps? (2) How does urbanisation alter community composition? (3) What roles do temperature, green cover, landscape diversity and fragmentation play in shaping these responses? (4) Do pollinator guilds (bees versus wasps) exhibit divergent responses to urban environmental conditions? By linking landscape structure to multiple ecological and fitness-related outcomes across cities, this study advances our understanding of urban evolutionary ecology and provides critical insights for biodiversity conservation in increasingly urbanised environments.

## 2. Materials and Methods

### 2.1 Study sites

To assess how urbanisation influences cavity-nesting Hymenopteran biodiversity, community composition and fitness, as well as the underlying environmental drivers, we launched the ‘Urbeevol Project’ a large citizen-science project in 2022 in Leipzig and Hamburg, Germany (Figure 1). Participants (n = 286; 178 in Hamburg, 108 in Leipzig; Figure 1) were selected to ensure spatially representative sampling along urbanisation gradients within each city. To ensure spatial independence, we maintained a minimum sampling distance of 500 m between participants, exceeding the typical foraging range of most insect pollinators (Gathmann & Tscharntke, 2002; Greenleaf et al., 2007).

**FIGURE 1.**
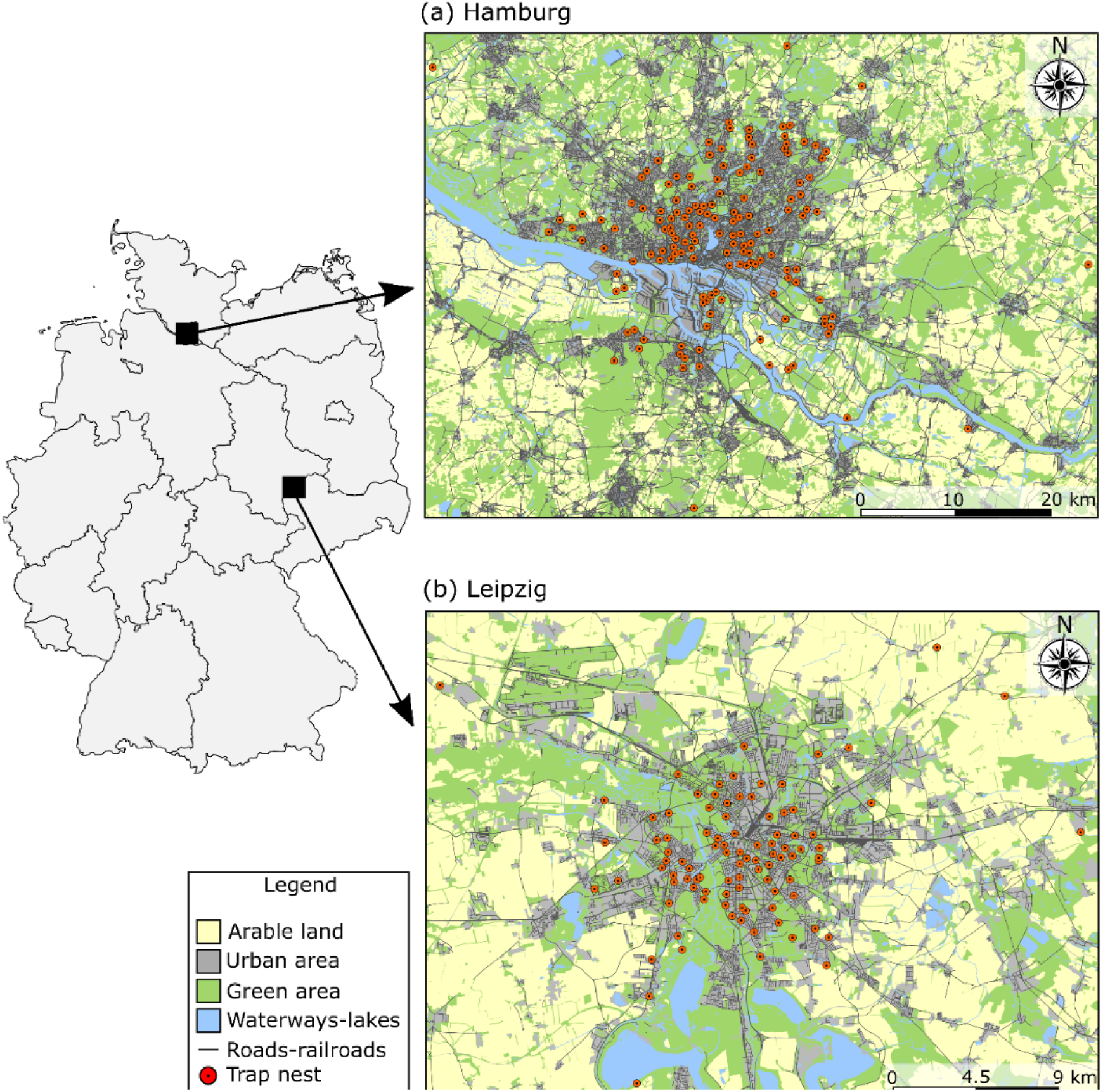
Sampling locations in **(a)** Hamburg and **(b)** Leipzig. Maps were created in R using OpenStreetMap data. Data licensed by OpenStreetMap Foundation (OSMF) under ODbL (https://opendatacommons.org/licenses/odbl/).

### 2.2 Sample collection, processing and species identification using COI barcoding

Participants installed trap nests on south-facing outdoor surfaces (e.g., balconies, carports, terraces) with high sun exposure and minimal shading (Everaars et al., 2011). Each trap nest consisted of a plastic tube (11 cm in diameter, 17 cm in length) randomly filled with 65-90 bamboo stems (15 cm length) with cavity diameters ranging from 3-10 mm. Trap nests were deployed in early March 2022 to accommodate species with varying phenologies. We verified correct placement using geotagged photographs submitted by participants.

In December 2022, trap nests (n = 251) were collected and overwintered outdoors at Martin-Luther University Halle-Wittenberg. In January 2023, occupied bamboo stems (n = 6,391) were transferred to cotton-sealed glass test tubes and incubated at room temperature to simulate spring emergence. Emerging individuals were collected in 100% ethanol, stored at -20 °C and counted as a proxy for reproductive output. At the end of the emergence period, nests were dissected to record the number of dead individuals. We identified bees and wasps to species using identification keys and DNA barcoding of the COI gene (Ratnasingham & Hebert, 2007) (Supplementary Methods 1). Bee and wasp specimens are kept at the General Zoology research group, Martin-Luther University Halle-Wittenberg. Sequences obtained from COI barcoding were submitted to the NCBI GenBank database and are available under Accession Numbers PV602798 - PV603255.

### 2.3 Species richness, occupancy, community composition and fitness

Species richness was defined as the total number of species recorded at each sampling site, representing community-level diversity. The number of nests referred to the total count of occupied stems, indicating the level of nesting activity. Reproductive output was measured as the number of successfully emerged offspring per occupied stem, reflecting reproductive success. Offspring survival was calculated as the proportion of provisioned brood that successfully emerged as adults, providing insight into post-provisioning survival rates. Demographic status was defined by reproductive output, measured as the number of adult offspring per individual nest. Nests with reproductive output ≥ 2 were classified as sources, while those with < 2 were classified as sinks.

To quantify community composition and dissimilarity across the urban gradient, we calculated pairwise beta diversity using the Jaccard index (presence-absence data), partitioned into turnover and nestedness-resultant components (Baselga, 2010, 2012) using the ‘betapart’ R package v. 1.6 (Baselga & Orme, 2012) (Supplementary Methods 2).

### 2.4 Environmental variables

Environmental drivers of biodiversity, community composition and fitness were assessed using a suite of landscape-scale variables within a 400 m buffer, approximating the foraging range of solitary Hymenoptera (Gathmann & Tscharntke, 2002; Zurbuchen et al., 2010). The proportion of impervious surfaces around each site was quantified using Quantum GIS v.

3.16.0 and data from CORINE Land Cover (European Environment Agency, 2019). The proportion of green cover (i.e., parks, allotment gardens, meadows, forests, cemeteries, scrublands), landscape diversity (Supplementary Methods 2) and landscape fragmentation (i.e. edge density; Supplementary Methods 2) were calculated in QGIS v3.16.0 using regional land use layers from Geofabrik GmbH, based on OpenStreetMap data. Additionally, the median land surface temperature (LST) was calculated from Landsat-9 satellite imagery (March–September 2022) following the approach by Ermida et al., (2020), with a minor modification to compute seasonal medians. Pairwise Pearson correlation coefficients were calculated among all environmental variables to characterise their interrelationships (Figure S1).

### 2.5 Statistical analyses

We used a Bayesian hierarchical modelling framework to assess the effects of urbanisation on cavity-nesting Hymenoptera. Specifically, to isolate the effects of urbanisation, we first implemented Bayesian generalised linear mixed models (GLMMs) using impervious surface cover as a single predictor on five response variables: (1) species richness, (2) number of nests, (3) reproductive output, (4) offspring survival and (5) demographic status. All analyses were conducted separately for bees and wasps. For offspring survival, we specified a binomial error structure with a logit link function, modelling the probability of surviving individuals per total number of individuals. For demographic status, we specified a Bernoulli error structure with a logit link function, modelling the probability of a nest functioning as a demographic source versus a sink. Species richness and number of nests were modelled using a Poisson error structure, while reproductive output was fitted using a negative binomial error structure to account for overdispersion, with an exponential link function. To account for hierarchical data structure, we included city as a random factor in species richness models, trap nests nested within cities for nest numbers and demographic status models, and stems nested within trap nests, nested within cities for reproductive output and offspring survival models. To address non-independence among species due to shared evolutionary history, pairwise evolutionary distances based on their phylogenies were transformed into a covariance matrix using the Ornstein–Uhlenbeck (L1) kernel (McElreath, 2016). This matrix was then included as a species-level random effect correlation structure in the number of nests, reproductive output, offspring survival and demographic status models. For the respective phylogenies, we constructed a neighbour-joining phylogeny for wasp species using COI sequences retrieved from NCBI GenBank following an alignment using the ClustalW algorithm (Thompson et al., 1994), employing the GTR+Γ substitution model (Tavaré, 1984; Yang, 1994) and incorporated a published phylogeny for bees (Henríquez-Piskulich et al., 2024).

To identify additional environmental drivers beyond urbanisation, we then repeated the analyses, replacing impervious surface cover with four ecologically relevant landscape-scale predictors: (1) landscape diversity (LD), (2) landscape fragmentation (LF), (3) median land surface temperature (LST) and (4) proportion of green cover (GC). All models retained the same error structures and random effect specifications described above.

To investigate how urbanisation and environmental variables affect community composition, we employed Bayesian GLMMs, using beta diversity and its components (turnover and nestedness) as response variables. The analyses were conducted separately for bees and wasps. We calculated absolute pairwise distances in impervious surface cover (|Δ_IMP_|) between sampling sites to model compositional dissimilarity along the urban gradient, while controlling for geographic distance (D_GEO_) as a proxy for spatial dispersal limitation. To further explore the influence of landscape structure, we modelled beta diversity and its components as functions of absolute differences in: (1) landscape diversity (|Δ_LD_|), (2) landscape fragmentation (|Δ_LF_|), (3) median land surface temperature (|Δ_LST_|) and (4) proportion of green cover (|Δ_GC_|) and (5) geographic distance (D_GEO_). All community dissimilarity models were fitted using an ordered beta distribution (Kubinec, 2023). To leverage the multi-city design, we first computed pairwise community comparisons within each city and then combined them into a joint dataset, including city as a random factor in the models.

All predictors were standardised (mean = 0, SD = 1) prior to analysis to ensure comparability of effect sizes. Models were implemented in R statistical software (R Core Team, 2024) using the ‘rstan’ package v. 2.36.2 (Guo et al., 2015) as an interface to Stan (Stan Development Team, 2024), employing the No-U-Turn Sampler (NUTS) (Hoffman & Gelman, 2014) for efficient posterior sampling. Model assumptions, convergence diagnostics and posterior predictive checks were evaluated using the ‘shinystan’ R package (Gabry & Veen, 2015; Gelman et al., 2014). We computed the probability of direction (pd) for each predictor using the ‘bayestestR’ R package (Makowski, Ben-Shachar, & Lüdecke, 2019) and followed Bayesian reporting standards (Makowski, Ben-Shachar, Chen, et al., 2019). All data handling and visualisation were conducted using the following R packages: datawizard, dplyr, HDInterval, tidyr, tidybayes, purr, ggplot2 and patchwork (Kay, 2018; Meredith & Kruschke, 2016; Patil et al., 2022; Pedersen, 2019; Wickham et al., 2014; Wickham & Henry, 2015).

## 3. Results

We deployed 286 trap nests across a wide range of urban, suburban and peri-urban environments in the cities of Hamburg and Leipzig, Germany, to capture variation in urbanisation intensity and landscape environmental structure (Figure 1). Of the 286 trap nests deployed, we successfully retrieved 251. In addition, 19 trap nests were excluded due to loss of site identity during transport or poor condition. This resulted in 232 trap nests comprising 6,391 occupied stems. From these, we identified 16 species of bees and 23 species of wasps (Tables S1 & S2). Across both cities, trap nests had an average occupancy of 27% (SD = 25%). Bee occupancy was at 22%, while wasp occupancy was at 5%. On average, bees produced three offspring per stem (SD = 3) with a survival rate of 74%, while wasps produced two offspring per stem (SD = 2) with a survival rate of 73%. 60% (SD = 49%) of bee nests and 57% (SD = 50%) of wasp nests functioned as demographic sources.

### 3.1 Effects of urbanisation on richness, number of nests, reproductive output, offspring survival and demographic status

Species richness declined with increasing impervious surface cover, with a stronger effect in wasps (bees: β_IMP_ = -0.112, 95% CI [-0.19, -0.03]; wasps: β_IMP_ = -0.244, 95% CI [-0.36, - 0.12]) (Figure 2, Figure S2, Tables S3 & S4). Number of nests did not show strong associations with impervious cover for either group (bees: β_IMP_ = -0.045, 95% CI [-0.13, 0.05], PD = 83.3%; wasps: β_IMP_ = -0.127, 95% CI [-0.28, 0.03], PD = 93.9%) (Figure 2, Figure S2, Tables S5 & S6). However, reproductive output declined with urbanisation in both guilds (bees: β_IMP_ = -0.076, 95% CI [-0.15, -0.01], PD = 98.4%; wasps: β_IMP_ = -0.112, 95% CI [-0.19, -0.03], PD = 99.7%) (Figure 2, Figure S2, Tables S7 & S8). Offspring survival also decreased with urbanisation, most steeply for bees (bees: β_IMP_ = -0.396, 95% CI [-0.60, -0.19], PD = 100%; wasps: β_IMP_ = -0.203, 95% CI [-0.39, -0.02], PD = 98.4%) (Figure 2, Figure S2, Tables S9 & S10). Urbanisation further reduced the likelihood of nests functioning as demographic source units in bees but not in wasps (bees: β_IMP_ = -0.127, 95% CI [-0.24, -0.02], PD = 98.9%; wasps: β_IMP_ = -0.0862, 95% CI [-0.24, 0.06], PD = 87.0%) (Figure 2, Figure S2, Tables S11 & S12).

**FIGURE 2.**
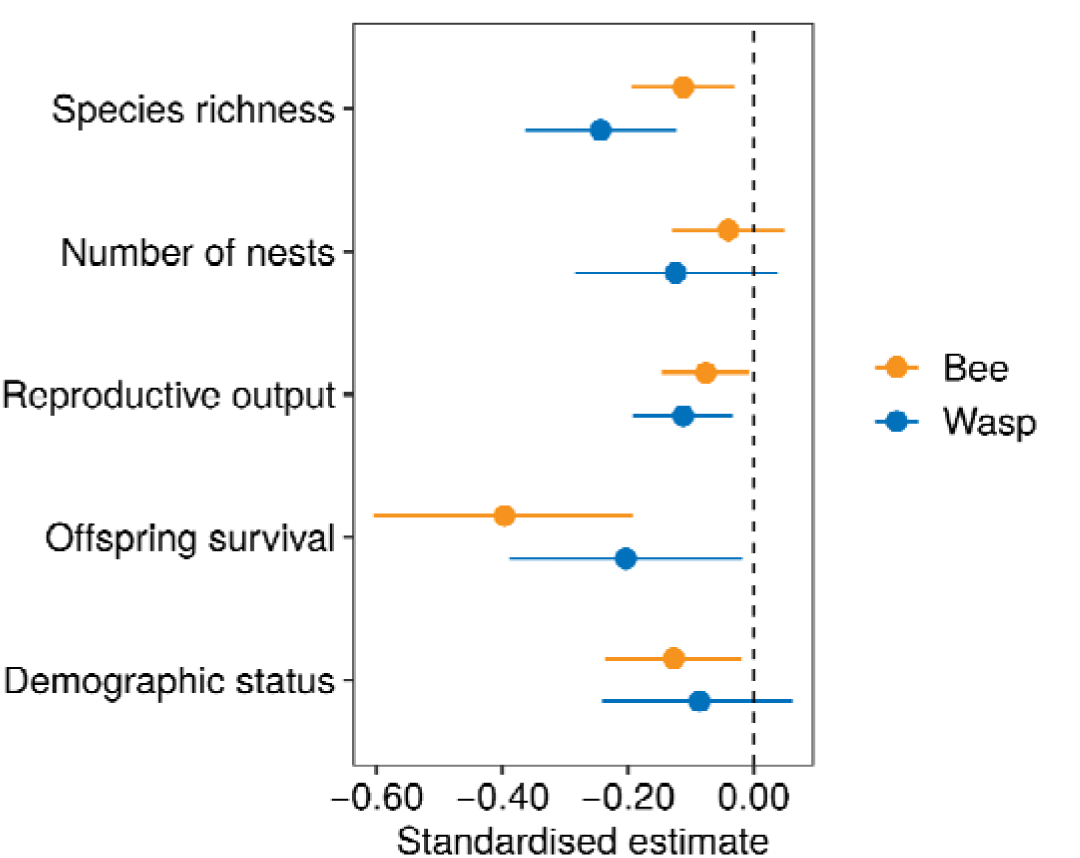
Standardised parameter estimates for impervious surface cover with 95% credible intervals across all five response variables (i.e., species richness, number of nests, reproductive output, offspring survival and demographic status).

### 3.2 Environmental drivers of bee and wasp responses

We conducted a more detailed analysis incorporating multiple landscape-scale variables, landscape diversity (LD), green cover (GC), land-surface temperature (LST) and landscape fragmentation (LF), to identify the key environmental drivers shaping cavity-nesting Hymenoptera responses to urbanisation (Figure 3; Tables S13-22).

**FIGURE 3.**
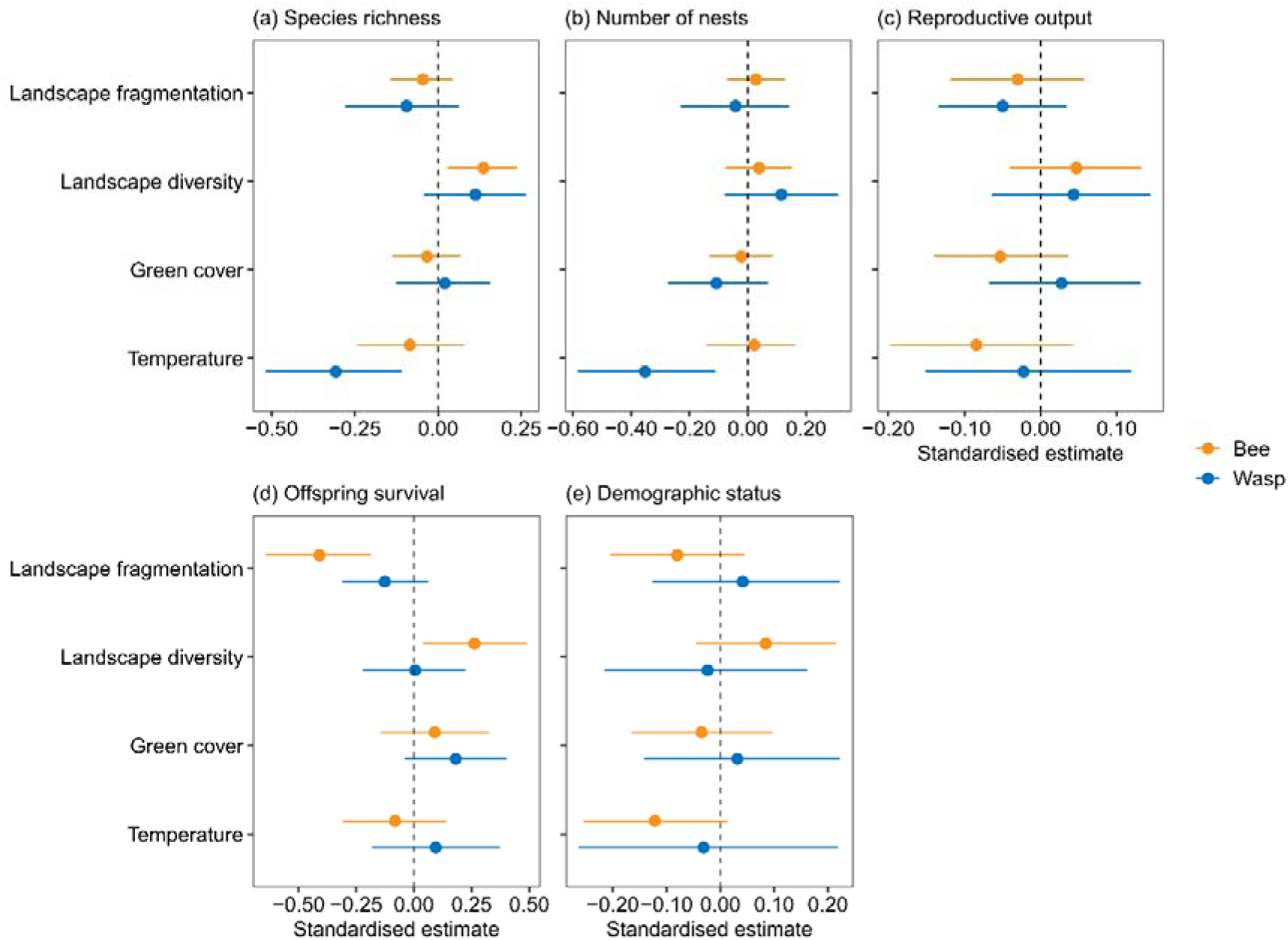
Standardised parameter estimates with 95% credible intervals from **(a)** species richness, **(b)** number of nests, **(c)** reproductive output, **(d)** offspring survival models and **(e)** demographic status models.

Bee species richness increased with greater landscape diversity (β_LD_ = 0.137, 95% CI [0.03, 0.24], PD = 99.4%; Figures 3a & 4a). In contrast, wasp species richness declined with increasing land-surface temperature (β_LST_ = -0.307, 95% CI [-0.52, -0.11], PD = 99.9%; Figures 3a & 4b). Other predictors showed no consistent effects on species richness (Figure 3a).

**FIGURE 4.**
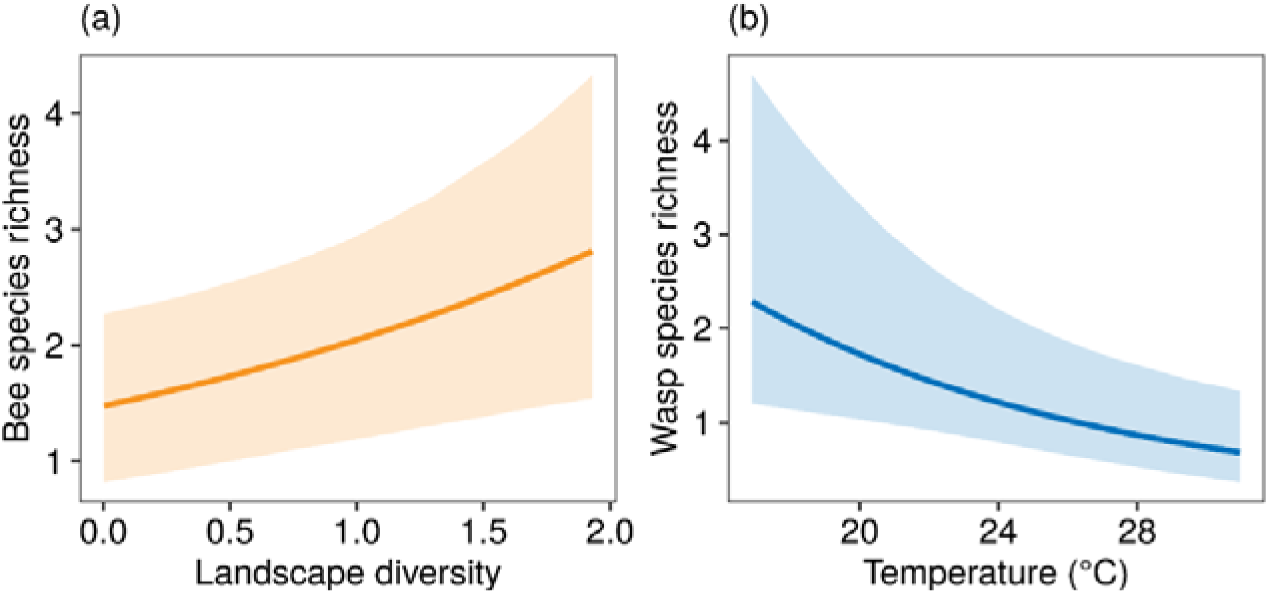
Relationships between **(a)** bee species richness and landscape diversity, **(b)** wasp species richness and land-surface temperature (°C). Lines represent the predicted relationships and shaded areas correspond to 95% credible intervals.

The number of bee nests showed no strong relationships with any landscape predictor (Figure 3b). The number of wasp nests, however, declined significantly with increasing land-surface temperature (β_LST_ = -0.354, 95% CI [-0.59, -0.11], PD = 99.8%; Figures 3b & 5). Other predictors showed no consistent effects on the number of wasp nests (Figure 3b).

**FIGURE 5.**
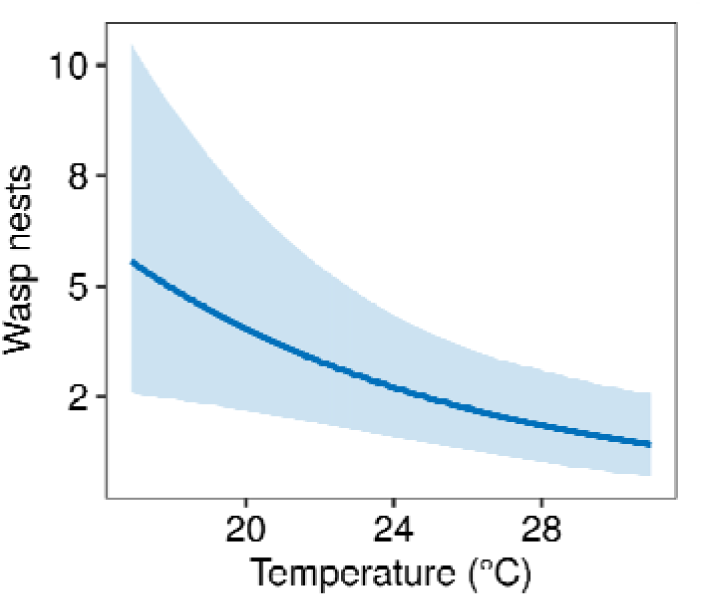
Relationship between the number of wasp nests and land-surface temperature (°C). Line represents the predicted relationship and shaded area corresponds to 95% credible interval.

No landscape variable was strongly associated with reproductive output in either guild (Figure 3c). However, bee offspring survival increased with landscape diversity (β_LD_ = 0.263, 95% CI [0.04, 0.49], PD = 99.0%; Figures 3d & 6a) and declined with increasing landscape fragmentation (β_LF_ = -0.409, 95% CI [-0.64, -0.19], PD = 100%; Figures 3d & 6b). Wasp offspring survival exhibited a weak positive trend with green cover (β_GC_ = 0.182, 95% CI [-0.04, 0.4], PD = 94.5%; Figures 3d & 6c). Other predictors showed no consistent effects on offspring survival (Figure 3d).

**FIGURE 6.**
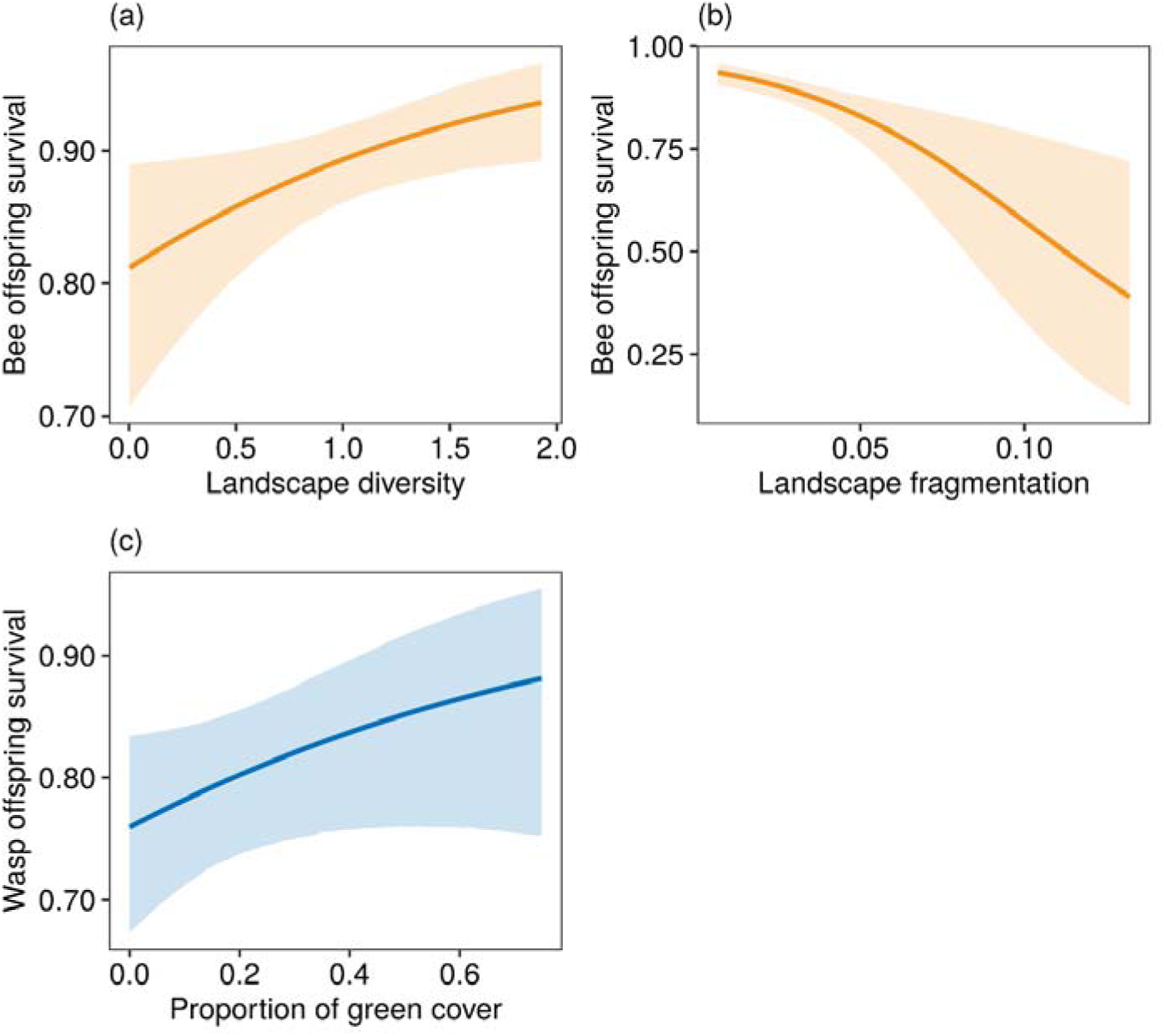
Relationships between bee offspring survival and **(a)** landscape diversity, **(b)** landscape fragmentation and **(c)** wasp offspring survival and proportion of green cover. Lines represent the predicted relationships and shaded areas correspond to 95% credible intervals.

Demographic source status in bees declined with increasing land-surface temperatures (β_LST_ = -0.122, 95% CI [-0.25, 0.01], PD = 95.9%; Figures 3e & 7). No landscape variable was strongly associated with demographic status in wasps (Figure 3e).

**FIGURE 7.**
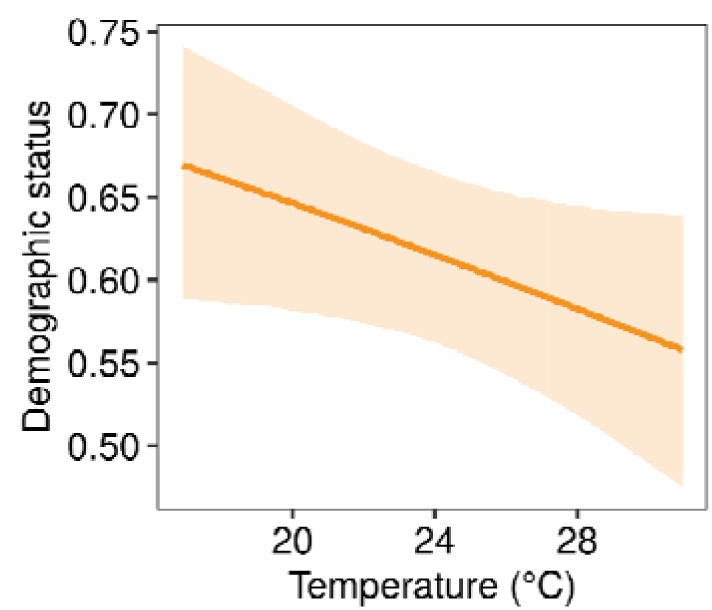
Relationships between bee demographic status and temperature. Line represents the predicted relationship and shaded area corresponds to 95% credible interval.

### 3.4 Effects of urbanisation on community composition

We used Bayesian models to relate pairwise beta diversity and its components (turnover and nestedness) to differences in impervious cover and geographic distance (Figure 8; Figure S3; Tables S23-28). Bee communities were more dissimilar (total beta diversity) across greater differences in impervious cover (β_|ΔIMP|_ = 0.039, 95% CI [0.02, 0.05], PD = 100%) and geographic distance (β_|DGEO|_ = 0.029, 95% CI [0.02, 0.04], PD = 100%) (Figure 8). Our models revealed increased species turnover (higher replacement) with increased differences in urbanisation (β_|ΔIMP|_ = 0.023, 95% CI [0.01, 0.04], PD = 99.6%), whereas nestedness remained unaffected (β_|ΔIMP|_ = -0.002, 95% CI [-0.02, 0.02], PD = 59.1%) (Figure 8). We furthermore found geographic distance to be negatively associated with turnover (β_|DGEO|_ = -0.035, 95% CI [-0.05, -0.02], PD = 100%) and positively associated with nestedness (β_|DGEO|_ = 0.043, 95% CI [0.02, 0.07], PD = 100%) (Figure 8). In contrast, wasp beta diversity and its components showed no credible association with urbanisation (Figure 8). Only nestedness increased with geographic distance (β_|DGEO|_ = 0.056, 95% CI [0.02, 0.1], PD = 99.7%; Figure 8).

**FIGURE 8.**
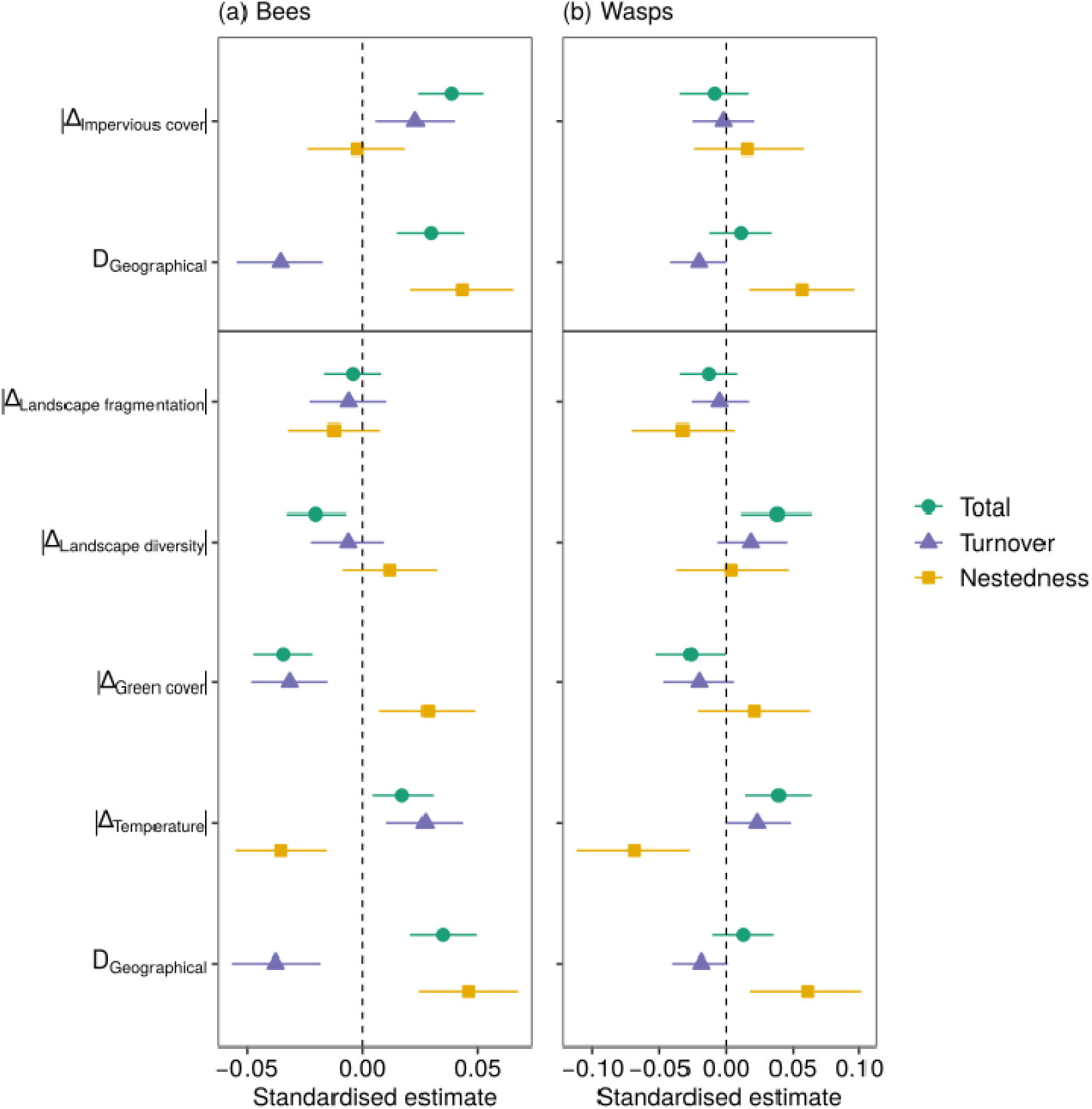
Standardised parameter estimates with 95% credible intervals from **(a)** bee and **(b)** wasp community composition models.

We next examined how absolute differences in environmental variables (i.e., landscape diversity, green cover, land-surface temperature and landscape fragmentation) shaped overall beta diversity, turnover and nestedness (Figure 8; Figure S4; Tables S29-34). For bees, total beta diversity increased with land-surface temperature differences (β_|ΔLST|_ = 0.017, 95% CI [0, 0.03], PD = 99.5%) and decreased with divergence in landscape diversity (β_|ΔLD|_ = -0.020, 95% CI [-0.03, -0.01], PD = 99.9%) and green cover (β_|ΔGC|_ = -0.034, 95% CI [-0.05, -0.02], PD = 100%) (Figure 8). Turnover increased with land-surface temperature differences (β_|ΔLST |_ = 0.027, 95% CI [0.01, 0.04], PD = 100%), but declined with divergence in green cover (β_|ΔGC|_ = -0.031, 95% CI [-0.05, -0.01], PD = 100%) (Figure 8). Nestedness increased with differences in green cover (β_|ΔGC|_ = 0.028, 95% CI [0.01, 0.05], PD = 99.6%) but declined with land-surface temperature differences (β_|ΔLST|_= -0.035, 95% CI [-0.05, -0.02], PD = 99.9%) (Figure 8). For wasps, total beta diversity increased with land-surface temperature differences (β_|ΔLST|_=0.039, 95% CI [0.01, 0.06], PD = 99.9%) and landscape diversity (β_|ΔLD|_=0.038, 95% CI [0.01, 0.06], PD = 99.8%), but declined with increasing differences in green cover (β_|ΔGC|_=-0.026, 95% CI [-0.05,0], PD = 97.8%) (Figure 8). Turnover increased with differences in land-surface temperature (β_|ΔLST|_ = 0.023, 95% CI [0, 0.05], PD = 97.2%), while other predictors were inconclusive (Figure 8). Nestedness declined with increasing differences in land-surface temperature (β_|ΔLST|_ = -0.068, 95% CI [-0.11, -0.03], PD = 99.9%) and increased with fragmentation differences (β_|ΔLF|_ = -0.032, 95% CI [-0.07, 0.01], PD = 95.3%) (Figure 8).

## 4. Discussion

### 4.1 Urbanisation reduces diversity and fitness while altering community composition

The ecological consequences of urbanisation for insects remain highly context-dependent, with studies reporting contrasting responses often shaped by life history traits, ecological flexibility and trophic position of the taxa studied (Banaszak-Cibicka & Żmihorski, 2012; Liang et al., 2023; Theodorou, Radzevičiūtė, et al., 2020). Cavity-nesting Hymenoptera are sometimes considered potential urban beneficiaries due to the abundance of artificial structures offering nesting cavities (Banaszak-Cibicka & Żmihorski, 2012; Liang et al., 2023; Wenzel et al., 2020). However, our findings demonstrate that urbanisation consistently exerts negative effects on these taxa, reducing species richness, reproductive output and offspring survival in both bees and wasps. Furthermore, our results indicate urbanisation reduced the likelihood of bee nests functioning as a demographic source. These patterns, observed across two cities and more than 230 trap nests, indicate that urban environments act as low-quality habitats for cavity-nesting Hymenoptera. The most pronounced declines were in offspring survival, particularly for bees, highlighting that nesting presence alone is an unreliable proxy for habitat quality. Despite relatively stable nest numbers across the cityscape, the decoupling of nesting activity from reproductive success and from the probability that nests function as demographic sources suggests the operation of ecological trap dynamics (Pulliam, 1988). Urban areas may attract nesting individuals while ultimately failing to support self-sustaining populations, highlighting the need to assess not only species richness and nest numbers but also reproductive output, offspring viability and demographic sustainability when evaluating habitat quality.

Our results also support the concept of urban filtering (Knop, 2016; McKinney, 2006; Rivest & Kharouba, 2024). Bee communities exhibited increasing beta diversity and species turnover with urbanisation, suggesting compositional shifts likely driven by differential persistence of disturbance-tolerant versus sensitive species. Interestingly, greater geographic distance between sites led to increased beta diversity in bees, primarily due to nestedness; indicating that species-poor communities are subsets of richer ones, rather than harbouring distinct species. This points to spatially structured species loss, likely reflecting dispersal limitation and the dominance of generalists in environmentally filtered urban systems. In contrast, cavity-nesting wasps show weaker spatial turnover, suggesting stronger trophic constraints and reduced dispersal or colonisation potential. This pattern suggests that urbanisation drives net species loss in wasps, likely due to their narrower ecological niches, higher trophic position and dependence on diverse prey resources (Ewers & Didham, 2006; Klaus et al., 2024; Moura et al., 2019; Powell & Taylor, 2017), which may be less available in urban settings (Theodorou, 2022). These guild-specific responses underscore the importance of considering ecological traits and trophic roles when assessing urban impacts. They also reveal the limitations of using species presence, abundance, or nest occupancy alone as indicators of habitat suitability. Although trap nests may appear well used in urban areas, our findings show that reproductive success and offspring survival can be substantially compromised, indicating a disconnect between habitat selection and quality. To explore the mechanisms underlying these patterns, the following sections assess the role of key environmental features on the fitness and community dynamics of bees and wasps across the urban gradient.

### 4.2 Landscape diversity as a buffer against urban stress for bees

Landscape diversity emerged as the strongest predictor of bee performance, positively associated with both richness and offspring survival. This finding aligns with a broad consensus that heterogeneous landscapes enhance pollinator diversity, likely by increasing resource availability and niche opportunities, reducing competition and facilitating species coexistence (Papanikolaou et al., 2017a; Senapathi et al., 2017). One likely mechanism underlying the positive effects of landscape diversity on offspring survival is nutritional: solitary bees rely on maternal pollen provisioning and diverse landscapes may offer a broader array of floral resources and pollen types that better meet the physiological needs of developing larvae (Filipiak et al., 2017; Parreño et al., 2024).

Landscape diversity also shaped community-level patterns. Bee communities became less compositionally distinct as differences in landscape diversity between sites increased, indicating community homogenisation across structurally divergent landscapes. This counterintuitive pattern suggests that increased landscape diversity does not necessarily support locally distinct or specialised bee assemblages. Instead, it may reflect the dominance of a core group of generalist species that are capable of exploiting a broad range of habitats, leading to similar community compositions even in ecologically dissimilar environments (Aronson et al., 2016; Bates et al., 2011; Deguines et al., 2016; Feng et al., 2024; Gathof et al., 2022). Such biotic homogenisation is well-documented in urban ecology, where environmental filtering favours species with broad ecological niches and high dispersal ability, while excluding specialists sensitive to habitat modification (Deguines et al., 2016; Gathof et al., 2022; Knop, 2016; Rivest & Kharouba, 2024). From a management perspective, these findings highlight the ecological value of spatial heterogeneity in urban environments. Enhancing landscape complexity can buffer bee populations against urban stressors, supporting the integration of diverse green spaces, including flowering streetscapes, community gardens, native plantings and structurally complex parks, into urban planning to foster resilient and functional pollinator communities.

Interestingly, the relationship between landscape diversity and community composition differed between the two pollinator guilds. While bees showed greater similarity in environments with high diversity differences, wasp communities became more dissimilar as landscape diversity differences increased. This contrast likely reflects fundamental differences in their trophic ecology. Wasps have specialised dietary needs that vary across life stages, including dependence on specific prey taxa for larval provisioning (Ewers & Didham, 2006; Klaus et al., 2024). In complex landscapes, the availability and distribution of prey may vary spatially, driving site-specific wasp assemblages. Conversely, generalist wasps may be less common in highly-diverse landscapes due to increased competition or predation risk. These guild-specific responses further highlight the importance of considering both resource type and trophic level when evaluating how environmental heterogeneity shapes urban insect communities.

### 4.3 Temperature as a limiting factor for Hymenoptera

Temperature is a major global determinant of Hymenoptera diversity (Orr et al., 2021), and in our study, land surface temperature emerged as one of the strongest environmental predictors of wasp responses. Both species richness and the number of wasp nests declined with increasing temperature, indicating heightened thermal sensitivity. In contrast, bees showed no clear response to temperature in terms of species richness and nest numbers, a pattern consistent with previously mixed findings (Geppert et al., 2023; Hamblin et al., 2017; Mayr et al., 2020; Papanikolaou et al., 2017b). However, our demographic status analysis revealed that bee nests are more likely to function as demographic sinks at warmer sites, suggesting that temperature impacts may primarily constrain long-term population viability rather than just nesting activity or abundance. Wasps, in contrast, showed no consistent association between temperature and demographic status, indicating different mechanisms of thermal vulnerability. This divergence, despite both groups being solitary and cavity-nesting, likely reflects underlying physiological, ecological and trophic differences. While neither group benefits from social thermoregulation, they may differ in thermal tolerance and behavioural plasticity. Some solitary bees, particularly those in urban areas, may exhibit broader thermal tolerance or flexible nesting phenology, enabling them to avoid extreme temperatures by adjusting nesting or foraging activity (Gonzalez et al., 2024; Maia-Silva et al., 2015; Willmer & Stone, 2004). Nonetheless, the increased prevalence of demographic sinks at high temperatures suggests that these behavioural strategies may be insufficient to buffer bees from thermal constraints on reproduction. Wasps may experience greater constraints in these adaptations (Käfer et al., 2012; O’Neill et al., 2023). Moreover, resource dependencies differ markedly.

Bee larvae are provisioned with pollen and nectar, resources that tend to be more reliably available in urban areas due to managed plantings (Baldock et al., 2019; Knapp et al., 2008), whereas wasps rely on live arthropod prey, with many species showing prey specialisation (Moura et al., 2019; Powell & Taylor, 2017). As temperatures rise, prey availability and diversity may decline (Cabon et al., 2024; Dampc et al., 2021), leading to resource-driven reproductive failure. Thus, wasps may be more exposed to indirect, bottom-up limitations, while bees appear directly thermally constrained in their ability to produce viable offspring, making them more likely to function as demographic sinks in warmer landscapes.

Community-level analyses support this interpretation. Differences in temperature were associated with increased dissimilarity and species turnover, and decreased nestedness, in wasps. This suggests that warming acts as a selective filter, excluding sensitive taxa and restructuring communities around a subset of tolerant species. The loss of nestedness indicates that communities in warmer sites host not only fewer species, but compositionally distinct assemblages. Bees, in contrast, showed no consistent response in richness, reproduction, or survival across the temperature gradient. However, demographic modelling revealed that nests in warmer areas functioned as population sinks, indicating insufficient reproduction to maintain local populations. Despite stable overall diversity, bee community composition still shifted with increasing temperature differences, indicating species turnover without net species loss. This likely reflects the replacement of thermally sensitive species by more heat-tolerant generalists, maintaining overall diversity while altering community structure. These results underscore that temperature shapes both fitness and community assembly, with stronger effects on predatory taxa. As cities continue to warm under climate change (Huang et al., 2019), thermally sensitive groups such as solitary wasps may face compounded risks from both habitat degradation and declining prey availability. These findings highlight the importance of preserving thermal refugia in urban landscapes. Interventions such as shade-providing vegetation, green roofs and microhabitat diversity can help mitigate urban heat and maintain cooler microclimates, offering crucial buffering for vulnerable Hymenoptera species. Understanding how temperature structures insect communities is particularly critical in urban systems, which may foreshadow broader ecological responses under future climate conditions (Huang et al., 2019).

### 4.4 Green cover and its functional role in supporting Hymenoptera communities

Contrary to expectations, we did not find a strong or consistent relationship between overall green cover and species richness in cavity-nesting Hymenoptera. This mirrors findings from previous studies that report mixed effects of green space extent on insect diversity (Ahrné et al., 2009; Anderson et al., 2023; Banaszak-Cibicka et al., 2016; Makinson et al., 2017). However, our fitness models revealed a weak positive trend between green cover and wasp survival, suggesting that green space might play a role in supporting wasp life histories, even if it does not directly influence species richness. This trend may reflect bottom-up constraints, particularly prey availability. Unlike bees, which can exploit floral resources across the cityscape, solitary wasps rely on sufficient and accessible prey for larval provisioning (Evans & West-Eberhard, 1970; Moura et al., 2019; Powell & Taylor, 2017; Witt, 1998). In areas with low green cover, prey populations may be sparse or require longer foraging trips. As shown in bumble bees (Pioltelli et al., 2024), increased foraging effort can expose nests to elevated risks of parasitism and predation, pressures that may be even more pronounced in wasps due to their prey specificity and solitary nesting behaviour. Thus, even modest reductions in green cover may impact wasp fitness.

While species richness was unaffected, greater differences in green cover between sites were associated with lower dissimilarity and species turnover and increased nestedness in Hymenoptera communities. This suggests that in more simplified landscapes, Hymenoptera assemblages become increasingly dominated by a core group of generalist species, leading to community homogenisation. Such patterns are consistent with habitat filtering, where only the most adaptable species persist under resource-limited environments (Deguines et al., 2016; Gathof et al., 2022; Knop, 2016; Rivest & Kharouba, 2024). Overall, our results indicate that green space quantity alone is an insufficient predictor of habitat quality. These findings highlight the importance of going beyond area-based assessments to consider the ecological functionality of urban green spaces. Ensuring that urban green areas are well-connected, managed appropriately, florally diverse and structurally complex may be essential for sustaining viable and diverse Hymenoptera communities in urban landscapes (Anderson et al., 2023; Banaszak-Cibicka et al., 2016; da Rocha-Filho et al., 2020; Turo & Gardiner, 2021).

### 4.5 Landscape fragmentation reduces bee survival

Among the environmental variables we examined, landscape fragmentation had the most pronounced negative effect on bee offspring survival. This aligns with a broad body of research showing that fragmentation reduces reproductive success in bees (Everaars et al., 2018; Nagamitsu et al., 2018) and other insects (Pichancourt et al., 2006). Fragmented landscapes typically consist of more isolated habitat patches, offering fewer floral and nesting resources and disrupting movement and gene flow (Couvet, 2002; Saccheri et al., 1998). For solitary bees, which have limited dispersal abilities, such isolation can increase inbreeding risk and reduce genetic diversity (Saccheri et al., 1998; Török et al., 2022; Williams et al., 2010), ultimately threatening population viability, particularly in rapidly changing urban environments (Johnson & Munshi-South, 2017). Fragmented habitats also tend to have higher edge-to-interior ratios, exposing nests to greater temperature fluctuations, desiccation and increased risks of predation and parasitism (Niebuhr et al., 2015; Stilley & Gabler, 2021). These abiotic and biotic stressors can reduce survival even when nesting opportunities are available. From a conservation standpoint, this underscores the need for urban planning strategies that prioritise habitat connectivity. Simply increasing the number of green spaces may be insufficient if those spaces remain ecologically isolated. Instead, efforts should focus on establishing well-connected, continuous habitat networks that reduce edge effects, promote gene flow and enhance population resilience.

### 4.6 Conclusions

Our study provides strong evidence that urbanisation affects cavity-nesting Hymenoptera through multiple ecological pathways, reducing diversity and reproductive fitness in bees and wasps. While overall patterns of species richness and reproductive output indicate negative impacts across taxa, only bees showed a shift in demographic status, becoming more likely to function as demographic sinks in warmer urban environments. These patterns are driven by distinct environmental variables, including temperature, green cover, landscape diversity and fragmentation that influence taxa in divergent ways. Effective conservation will require integrated, taxon-specific strategies that go beyond expanding green space, focusing instead on improving habitat quality, connectivity and thermal heterogeneity. As cities increasingly mirror future climate scenarios, they also offer valuable systems for studying biodiversity under rapid environmental change. To sustain resilient pollinator communities, we advocate for continued long-term, multi-city research, particularly studies that integrate ecological monitoring with fitness-based metrics such as reproductive success, offspring viability and demographic stability. These insights are essential for informing conservation policy aimed at supporting the persistence and ecological functionality of pollinator guilds in an urbanising world.

## Acknowledgements

We would like to thank our student helpers, Simon Dubielzig, Emil Cyranka, Tom Lauritz Hügel and Hannah Wachsmuth, for their valuable assistance. We are grateful to Dr. Hannes Hoffmann (BUKEA Hamburg) and Maik Hausotte (AfU Leipzig) for providing sampling permits.

## CRediT authorship contribution statement

Atilla Çelikgil: Conceptualisation, Data Curation, Formal Analysis, Investigation, Methodology, Writing – original draft, Writing – review & editing; Lars Henzschel: Investigation; Jonathan Sänger: Investigation; Pascal Dennis Krause: Investigation; Adrian Nemetschek: Investigation; Martin Husemann: Conceptualisation, Writing – review & editing; Panagiotis Theodorou. Conceptualisation, Funding Acquisition, Methodology, Project Administration, Resources, Supervision, Writing – original draft, Writing – review & editing

