## Supplementary Material for "Urbanisation drives biodiversity loss, fitness decline and community shifts in cavity-nesting Hymenoptera"

[**Supplementary Figure 4.** Relationships between differences in environmental conditions and components of bee and wasp β-diversity. Panels show the effects of differences in landscape diversity on **(a)** bee and **(b)** wasp β-diversity; differences in landscape fragmentation on **(c)** wasp nestedness; differences in green cover on **(d)** bee β-diversity, **(e)** bee turnover, **(f)** bee nestedness, and **(g)** wasp β-diversity; and differences in temperature on **(h)** bee β-diversity, **(i)** bee turnover, **(j)** bee nestedness, **(k)** wasp β-diversity, **(l)** wasp turnover, and **(m)** wasp nestedness. Lines represent predicted relationships and shaded areas correspond to 95% credible interval.	7](#_Toc204512914)

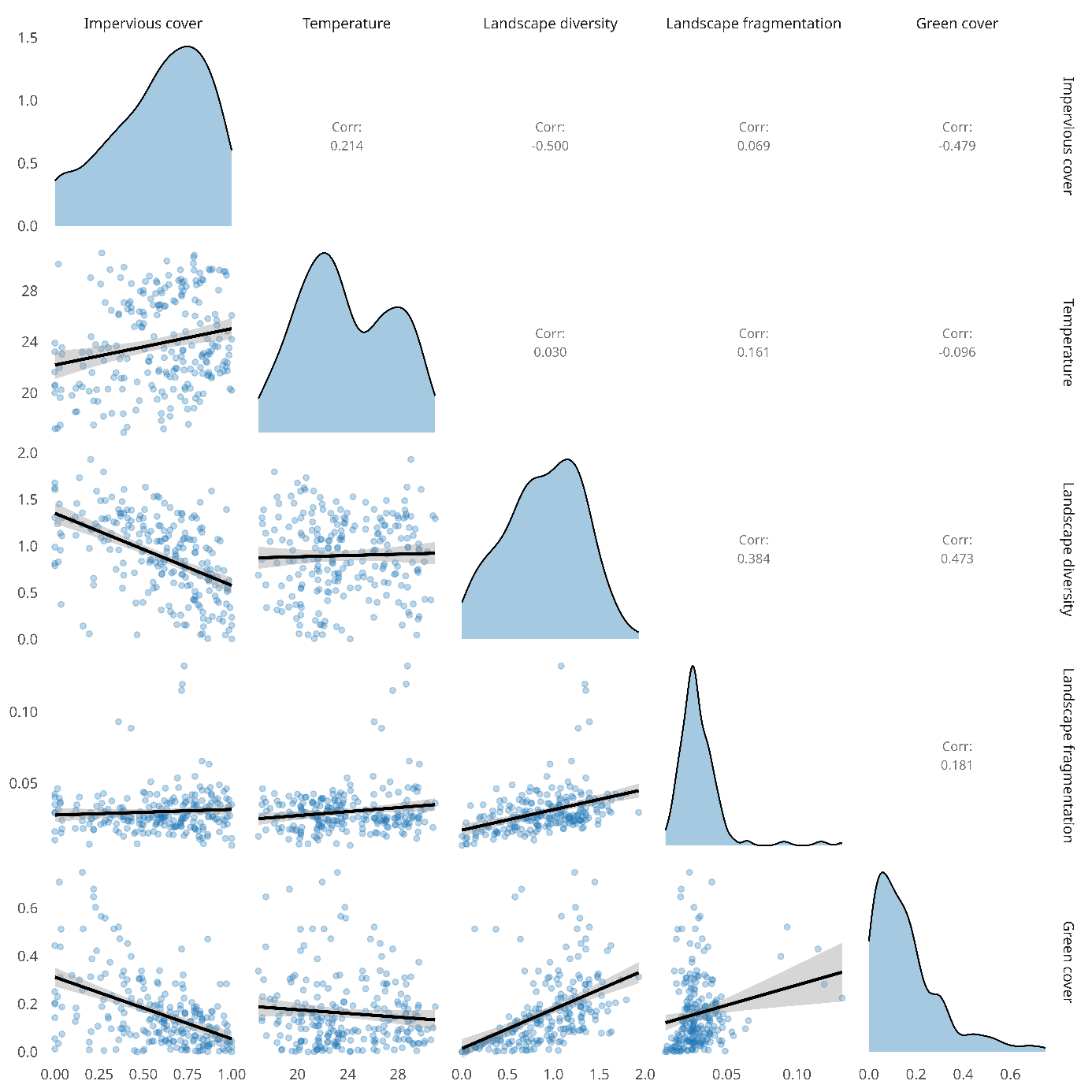

### **Supplementary Figure 1.** Pairwise relationships among five landscape variables: impervious cover, temperature, landscape diversity, landscape fragmentation and green cover. The lower triangle shows scatterplots with fitted linear regression lines and 95% confidence intervals. Diagonal panels display kernel density plots of each variable’s distribution. The upper triangle reports Pearson correlation coefficients.

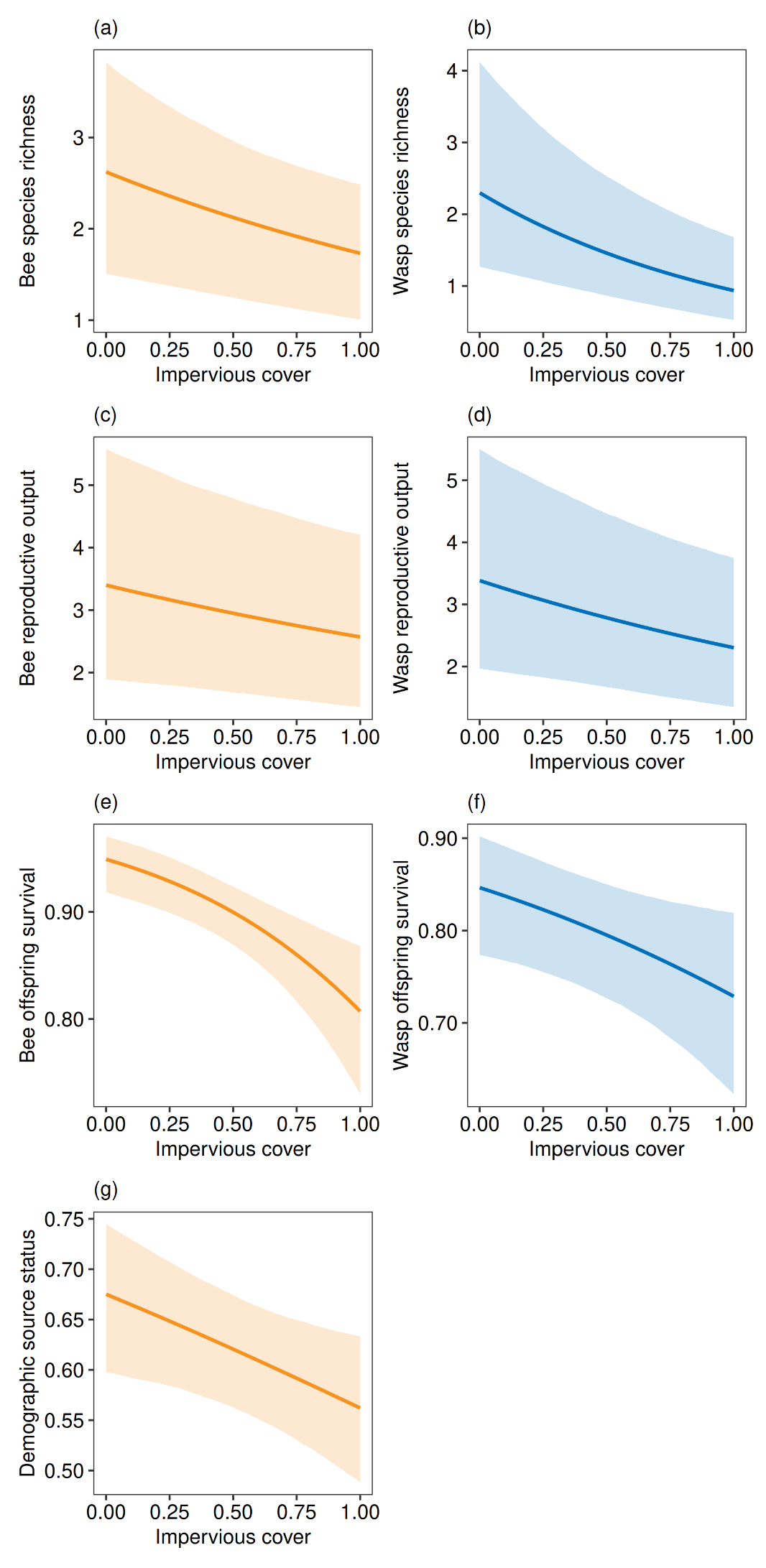

### **Supplementary Figure 2.** Relationships between impervious cover and (a) bee richness, (b) wasp richness, (c) bee abundance, (d) wasp abundance, (e) bee survival, (f) wasp survival, and (g) bee nest demographic source status. Lines represent the predicted relationships and shaded areas correspond to 95% credible interval.

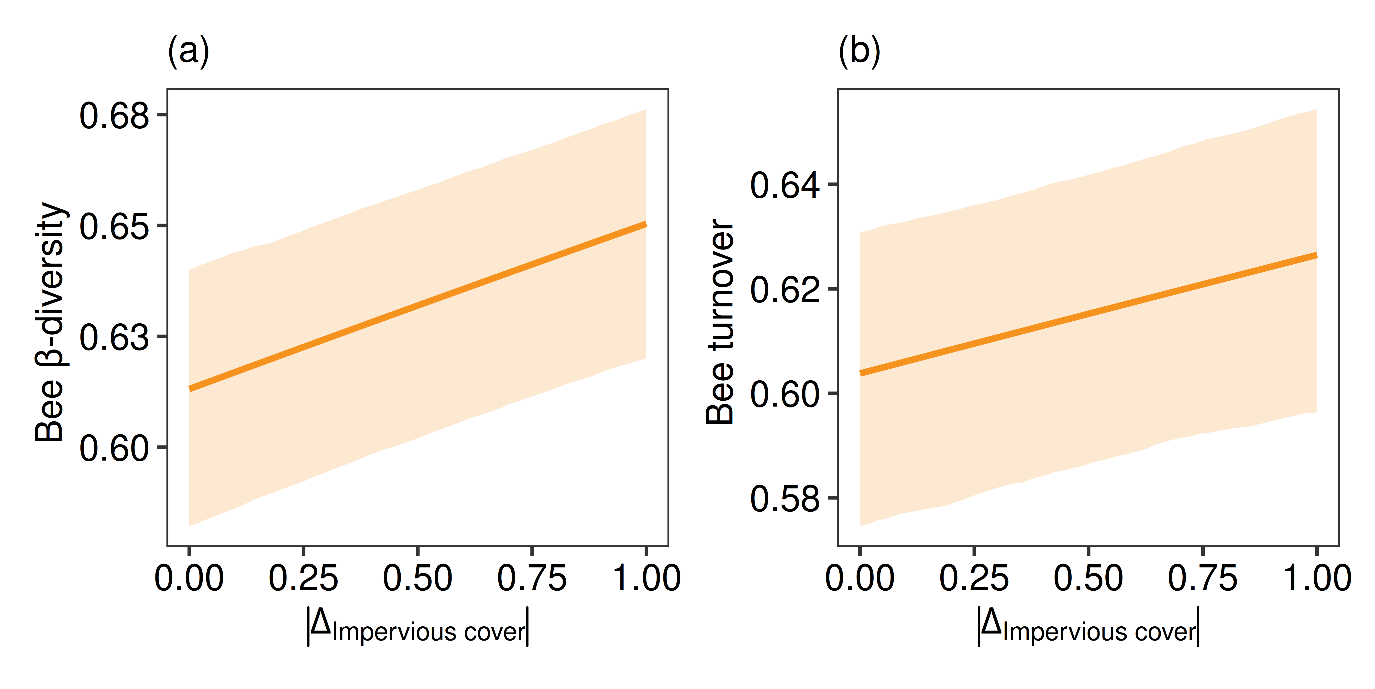

### **Supplementary Figure 3.** Relationships between impervious cover distance and bee (a) beta diversity and (b) turnover. Lines represent the predicted relationships and shaded areas correspond to 95% credible interval.

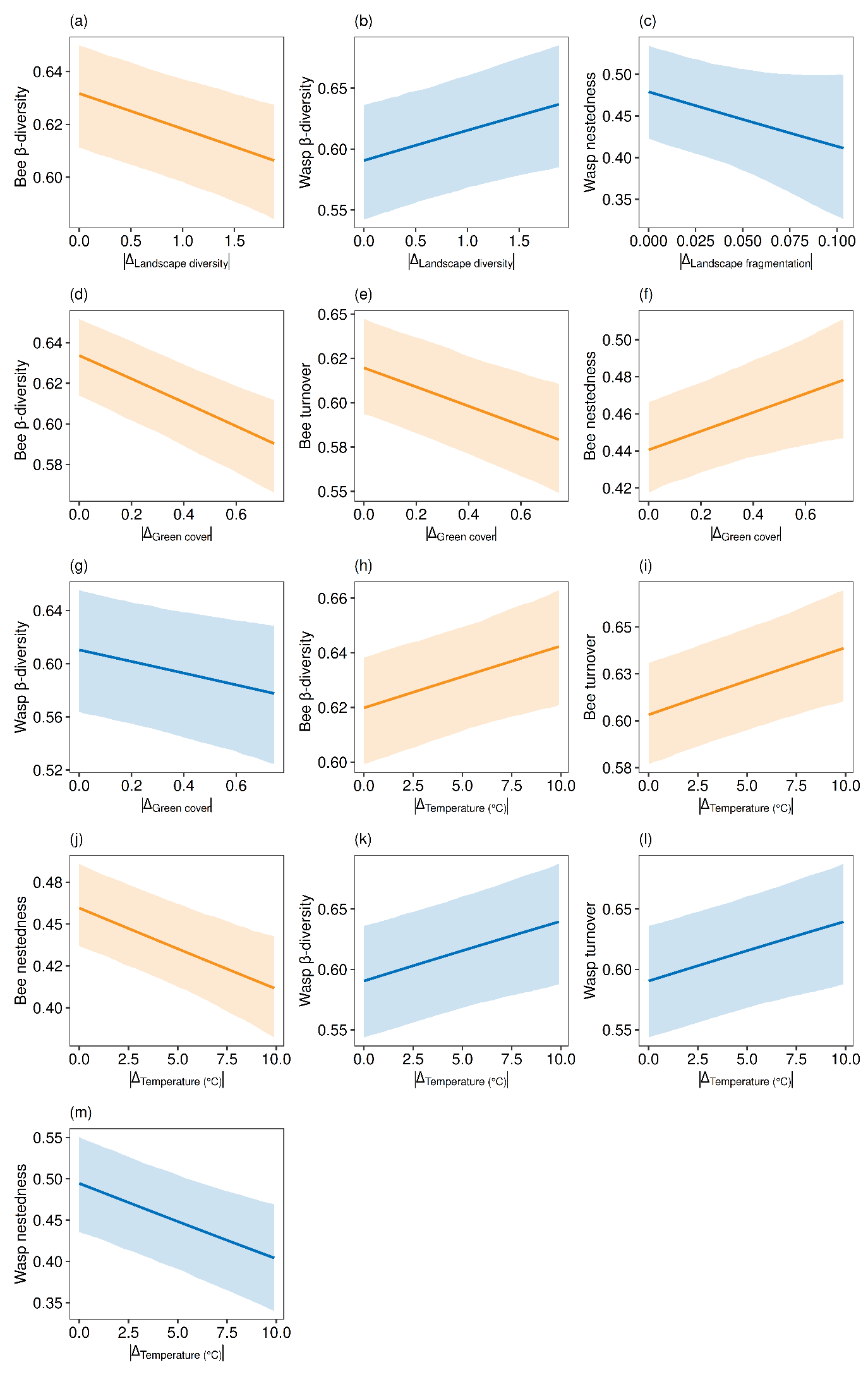

### **Supplementary Figure 4.** Relationships between differences in environmental conditions and components of bee and wasp β-diversity. Panels show the effects of differences in landscape diversity on **(a)** bee and **(b)** wasp β-diversity; differences in landscape fragmentation on **(c)** wasp nestedness; differences in green cover on **(d)** bee β-diversity, **(e)** bee turnover, **(f)** bee nestedness, and **(g)** wasp β-diversity; and differences in temperature on **(h)** bee β-diversity, **(i)** bee turnover, **(j)** bee nestedness, **(k)** wasp β-diversity, **(l)** wasp turnover, and **(m)** wasp nestedness. Lines represent predicted relationships and shaded areas correspond to 95% credible interval.

### **Supplementary Table 1.** Presence of bee species in trap nests across Leipzig and Hamburg.

| **Species** | **Leipzig** | **Hamburg** |
| --- | --- | --- |
| *Anthidium manicatum* | 1 | 0 |
| *Chelostoma rapunculi* | 4 | 19 |
| *Heriades truncorum* | 9 | 38 |
| *Osmia adunca* | 1 | 2 |
| *Hylaeus communis* | 29 | 79 |
| *Megachile alpicola* | 1 | 0 |
| *Megachile centuncularis* | 9 | 3 |
| *Megachile ericetorum* | 1 | 0 |
| *Megachile rotundata* | 19 | 7 |
| *Megachile versicolor* | 0 | 1 |
| *Osmia bicornis* | 70 | 75 |
| *Osmia brevicornis* | 16 | 3 |
| *Osmia caerulescens* | 24 | 22 |
| *Osmia cornuta* | 59 | 31 |
| *Osmia leaiana* | 2 | 2 |
| *Pseudoanthidium scapulare* | 4 | 0 |

### **Supplementary Table 2.** Presence of wasp species in trap nests across Leipzig and Hamburg.

| **Species** | **Leipzig** | **Hamburg** |
| --- | --- | --- |
| *Allodynerus rossii* | 0 | 1 |
| *Ancistrocerus antilope* | 11 | 4 |
| *Ancistrocerus gazella* | 7 | 17 |
| *Ancistrocerus nigricornis* | 2 | 9 |
| *Ancistrocerus trifasciatus* | 1 | 9 |
| *Auplopus carbonarius* | 0 | 1 |
| *Dipogon bifasciatus* | 1 | 0 |
| *Dipogon subintermedius* | 1 | 2 |
| *Discoelius dufourii* | 0 | 1 |
| *Euodynerus notatus* | 0 | 6 |
| *Passaloecus corniger* | 0 | 3 |
| *Passaloecus insignis* | 0 | 2 |
| *Pemphredon montana* | 2 | 2 |
| *Psenulus concolor* | 0 | 1 |
| *Psenulus fuscipennis* | 1 | 2 |
| *Psenulus pallipes* | 4 | 13 |
| *Symmorphus bifasciatus* | 1 | 8 |
| *Symmorphus crassicornis* | 2 | 16 |
| *Symmorphus gracilis* | 0 | 2 |
| *Symmorphus murarius* | 0 | 3 |
| *Trypoxylon clavicerum* | 3 | 2 |
| *Trypoxylon figulus* | 28 | 41 |
| *Trypoxylon medium* | 2 | 7 |

### **Supplementary Table 3.** Summary of Bayesian model examining the effect of urbanisation (impervious surface cover) on bee species richness.

| **Parameter** | **Median** | **Probability of Direction (%)** | **Effective Sample Size (ESS)** | **Rhat** | **90% CI** | **95% CI** |
| --- | --- | --- | --- | --- | --- | --- |
| alpha | 0.732 | 99.4 | 4,114 | 1.00 | [0.29, 0.99] | [0.18, 1.05] |
| beta_impervious surface cover | −0.112 | 99.6 | 9,490 | 1.00 | [-0.18, -0.04] | [-0.19, -0.03] |

### **Supplementary Table 4.** Summary of Bayesian model examining the effect of urbanisation (impervious surface cover) on wasp species richness.

| **Parameter** | **Median** | **Probability of Direction (%)** | **Effective Sample Size (ESS)** | **Rhat** | **90% CI** | **95% CI** |
| --- | --- | --- | --- | --- | --- | --- |
| alpha | 0.236 | 80.7 | 5,051 | 1.00 | [-0.16, 0.75] | [-0.22, 0.85] |
| beta_impervious surface cover | −0.244 | 100 | 8,107 | 1.00 | [-0.34, -0.14] | [-0.36, -0.12] |

### **Supplementary Table 5.** Summary of Bayesian model examining the effect of urbanisation (impervious surface cover) on the number of bee nests.

| **Parameter** | **Median** | **Probability of Direction (%)** | **Effective Sample Size (ESS)** | **Rhat** | **90% CI** | **95% CI** |
| --- | --- | --- | --- | --- | --- | --- |
| alpha | 1.04 | 100 | 11,908 | 1.00 | [0.58, 1.49] | [0.49, 1.58] |
| beta_impervious surface cover | −0.0435 | 83.3 | 13,334 | 1.00 | [-0.12, 0.03] | [-0.13, 0.05] |

### **Supplementary Table 6.** Summary of Bayesian model examining the effect of urbanisation (impervious surface cover) on the number of wasp nests.

| **Parameter** | **Median** | **Probability of Direction (%)** | **Effective Sample Size (ESS)** | **Rhat** | **90% CI** | **95% CI** |
| --- | --- | --- | --- | --- | --- | --- |
| alpha | 0.930 | 100 | 6,873 | 1.00 | [0.54, 1.33] | [0.46, 1.41] |
| beta_impervious surface cover | −0.127 | 93.9 | 9,134 | 1.00 | [-0.26, 0.01] | [-0.28, 0.03] |

### **Supplementary Table 7.** Summary of Bayesian model assessing the effect of urbanisation (impervious surface cover) on bee reproductive output.

| **Parameter** | **Median** | **Probability of Direction (%)** | **Effective Sample Size (ESS)** | **Rhat** | **90% CI** | **95% CI** |
| --- | --- | --- | --- | --- | --- | --- |
| alpha | 1.04 | 100 | 10,028 | 1.00 | [0.59, 1.46] | [0.51, 1.55] |
| beta_impervious surface cover | −0.0761 | 98.4 | 3,544 | 1.00 | [-0.14, -0.02] | [-0.15, -0.01] |

### **Supplementary Table 8.** Summary of Bayesian model assessing the effect of urbanisation (impervious surface cover) on wasp reproductive output.

| **Parameter** | **Median** | **Probability of Direction (%)** | **Effective Sample Size (ESS)** | **Rhat** | **90% CI** | **95% CI** |
| --- | --- | --- | --- | --- | --- | --- |
| alpha | 0.983 | 100 | 11,909 | 1.00 | [0.58, 1.41] | [0.51, 1.49] |
| beta_impervious surface cover | −0.112 | 99.7 | 12,813 | 1.00 | [-0.18, -0.05] | [-0.19, -0.03] |

### **Supplementary Table 9.** Summary of Bayesian model assessing the effect of urbanisation (impervious surface cover) on bee offspring survival.

| **Parameter** | **Median** | **Probability of Direction (%)** | **Effective Sample Size (ESS)** | **Rhat** | **90% CI** | **95% CI** |
| --- | --- | --- | --- | --- | --- | --- |
| alpha | 2.14 | 100 | 8,241 | 1.00 | [1.89, 2.38] | [1.83, 2.42] |
| beta_impervious surface cover | −0.396 | 100 | 6,889 | 1.00 | [-0.57, -0.23] | [-0.6, -0.19] |

### **Supplementary Table 10.** Summary of Bayesian model assessing the effect of urbanisation (impervious surface cover) on wasp offspring survival.

| **Parameter** | **Median** | **Probability of Direction (%)** | **Effective Sample Size (ESS)** | **Rhat** | **90% CI** | **95% CI** |
| --- | --- | --- | --- | --- | --- | --- |
| alpha | 1.40 | 100 | 6,554 | 1.00 | [1.08, 1.71] | [1.01, 1.77] |
| beta_impervious surface cover | −0.203 | 98.4 | 7,418 | 1.00 | [-0.36, -0.05] | [-0.39, -0.02] |

### **Supplementary Table 11.** Summary of Bayesian model assessing the effect of urbanisation (impervious surface cover) on bee nest demographic status.

| **Parameter** | **Median** | **Probability of Direction (%)** | **Effective Sample Size (ESS)** | **Rhat** | **90% CI** | **95% CI** |
| --- | --- | --- | --- | --- | --- | --- |
| alpha | 0.472 | 99.9 | 3,787 | 1.00 | [0.28, 0.67] | [0.23, 0.70] |
| beta_impervious surface cover | −0.127 | 98,9 | 3,406 | 1.00 | [-0.22, -0.04] | [-0.24, -0.02] |

### **Supplementary Table 12.** Summary of Bayesian model assessing the effect of urbanisation (impervious surface cover) on wasp nest demographic status.

| **Parameter** | **Median** | **Probability of Direction (%)** | **Effective Sample Size (ESS)** | **Rhat** | **90% CI** | **95% CI** |
| --- | --- | --- | --- | --- | --- | --- |
| alpha | 0.320 | 97.4 | 3,679 | 1.00 | [0.05, 0.60] | [0.00, 0.65] |
| beta_impervious surface cover | −0.0862 | 87.0 | 6,154 | 1.00 | [-0.22, 0.04] | [-0.24, 0.06] |

### **Supplementary Table 13.** Summary of Bayesian model evaluating the influence of urban environmental factors on bee species richness.

| **Parameter** | **Median** | **Probability of Direction (%)** | **Effective Sample Size (ESS)** | **Rhat** | **90% CI** | **95% CI** |
| --- | --- | --- | --- | --- | --- | --- |
| alpha | 0.706 | 99.3 | 6,584 | 1.00 | [0.25, 1] | [0.16, 1.06] |
| beta_green cover | −0.0330 | 74.1 | 9,358 | 1.00 | [-0.12, 0.05] | [-0.14, 0.07] |
| beta_ land surface temperature | −0.0848 | 84.1 | 7,824 | 1.00 | [-0.22, 0.05] | [-0.24, 0.08] |
| beta_ landscape diversity | 0.137 | 99.4 | 10,563 | 1.00 | [0.05, 0.22] | [0.03, 0.24] |
| beta_ edge density | −0.0453 | 83.1 | 12,332 | 1.00 | [-0.13, 0.03] | [-0.14, 0.04] |

### **Supplementary Table 14.** Summary of Bayesian model evaluating the influence of urban environmental factors on wasp species richness.

| **Parameter** | **Median** | **Probability of Direction (%)** | **Effective Sample Size (ESS)** | **Rhat** | **90% CI** | **95% CI** |
| --- | --- | --- | --- | --- | --- | --- |
| alpha | 0.0913 | 63.4 | 4,125 | 1.00 | [-0.2, 0.67] | [-0.24, 0.79] |
| beta_green cover | 0.0203 | 61.3 | 10,427 | 1.00 | [-0.1, 0.13] | [-0.13, 0.16] |
| beta_ land surface temperature | −0.307 | 99.9 | 9,402 | 1.00 | [-0.48, -0.14] | [-0.52, -0.11] |
| beta_ landscape diversity | 0.112 | 92.2 | 10,470 | 1.00 | [-0.02, 0.24] | [-0.04, 0.26] |
| beta_ edge density | −0.0947 | 87.1 | 11,017 | 1.00 | [-0.25, 0.04] | [-0.28, 0.06] |

### **Supplementary Table 15.** Summary of Bayesian model evaluating the influence of urban environmental factors on the number of bee nests.

| **Parameter** | **Median** | **Probability of Direction (%)** | **Effective Sample Size (ESS)** | **Rhat** | **90% CI** | **95% CI** |
| --- | --- | --- | --- | --- | --- | --- |
| alpha | 1.04 | 100 | 14,777 | 1.00 | [0.57, 1.5] | [0.48, 1.59] |
| beta_green cover | −0.0227 | 65.8 | 11,928 | 1.00 | [-0.11, 0.07] | [-0.13, 0.09] |
| beta_ land surface temperature | 0.0207 | 61.0 | 10,967 | 1.00 | [-0.12, 0.14] | [-0.14, 0.16] |
| beta_ landscape diversity | 0.0407 | 76.4 | 12,828 | 1.00 | [-0.05, 0.14] | [-0.07, 0.16] |
| beta_ edge density | 0.0263 | 70.3 | 13,810 | 1.00 | [-0.06, 0.11] | [-0.07, 0.13] |

### **Supplementary Table 16.** Summary of Bayesian model evaluating the influence of urban environmental factors on the number of wasp nests.

| **Parameter** | **Median** | **Probability of Direction (%)** | **Effective Sample Size (ESS)** | **Rhat** | **90% CI** | **95% CI** |
| --- | --- | --- | --- | --- | --- | --- |
| alpha | 0.972 | 100 | 11,482 | 1.00 | [0.55, 1.41] | [0.46, 1.5] |
| beta_green cover | −0.105 | 88.3 | 9,852 | 1.00 | [-0.25, 0.04] | [-0.28, 0.07] |
| beta_ land surface temperature | −0.354 | 99.8 | 5,893 | 1.00 | [-0.55, -0.15] | [-0.59, -0.11] |
| beta_ landscape diversity | 0.116 | 87.6 | 10,431 | 1.00 | [-0.05, 0.28] | [-0.08, 0.32] |
| beta_ edge density | −0.0451 | 68.2 | 12,185 | 1.00 | [-0.2, 0.11] | [-0.23, 0.14] |

### **Supplementary Table 17.** Summary of Bayesian model evaluating the influence of urban environmental factors on bee reproductive output.

| **Parameter** | **Median** | **Probability of Direction (%)** | **Effective Sample Size (ESS)** | **Rhat** | **90% CI** | **95% CI** |
| --- | --- | --- | --- | --- | --- | --- |
| alpha | 1.02 | 100 | 11,121 | 1.00 | [0.59, 1.46] | [0.5, 1.54] |
| beta_green cover | −0.0530 | 87.1 | 4,075 | 1.00 | [-0.13, 0.02] | [-0.14, 0.04] |
| beta_ land surface temperature | −0.0846 | 90.8 | 4,592 | 1.00 | [-0.18, 0.02] | [-0.2, 0.04] |
| beta_ landscape diversity | 0.0470 | 85.6 | 4,890 | 1.00 | [-0.03, 0.12] | [-0.04, 0.13] |
| beta_ edge density | −0.0300 | 75.5 | 4,698 | 1.00 | [-0.1, 0.04] | [-0.12, 0.06] |

### **Supplementary Table 18.** Summary of Bayesian model evaluating the influence of urban environmental factors on wasp reproductive output.

| **Parameter** | **Median** | **Probability of Direction (%)** | **Effective Sample Size (ESS)** | **Rhat** | **90% CI** | **95% CI** |
| --- | --- | --- | --- | --- | --- | --- |
| alpha | 0.982 | 100 | 7,761 | 1.00 | [0.58, 1.41] | [0.49, 1.49] |
| beta_green cover | 0.0275 | 71.1 | 5,839 | 1.00 | [-0.05, 0.11] | [-0.07, 0.13] |
| beta_ land surface temperature | −0.0223 | 62.7 | 5,085 | 1.00 | [-0.13, 0.09] | [-0.15, 0.12] |
| beta_ landscape diversity | 0.0432 | 79.0 | 7,067 | 1.00 | [-0.05, 0.13] | [-0.06, 0.14] |
| beta_ edge density | −0.0497 | 87.7 | 9,038 | 1.00 | [-0.12, 0.02] | [-0.13, 0.03] |

### **Supplementary Table 19.** Summary of Bayesian model evaluating the influence of urban environmental factors on bee offspring survival.

| **Parameter** | **Median** | **Probability of Direction (%)** | **Effective Sample Size (ESS)** | **Rhat** | **90% CI** | **95% CI** |
| --- | --- | --- | --- | --- | --- | --- |
| alpha | 2.11 | 100 | 7,589 | 1.00 | [1.85, 2.34] | [1.79, 2.39] |
| beta_green cover | 0.0908 | 77.5 | 6,635 | 1.00 | [-0.1, 0.28] | [-0.14, 0.33] |
| beta_ land surface temperature | −0.0814 | 76.5 | 7,183 | 1.00 | [-0.27, 0.1] | [-0.31, 0.14] |
| beta_ landscape diversity | 0.263 | 99.0 | 7,461 | 1.00 | [0.08, 0.45] | [0.04, 0.49] |
| beta_ edge density | −0.409 | 100 | 7,296 | 1.00 | [-0.6, -0.22] | [-0.64, -0.19] |

### **Supplementary Table 20.** Summary of Bayesian model evaluating the influence of urban environmental factors on wasp offspring survival.

| **Parameter** | **Median** | **Probability of Direction (%)** | **Effective Sample Size (ESS)** | **Rhat** | **90% CI** | **95% CI** |
| --- | --- | --- | --- | --- | --- | --- |
| alpha | 1.36 | 100 | 5,285 | 1.00 | [1.06, 1.67] | [0.99, 1.73] |
| beta_green cover | 0.182 | 94.5 | 5,177 | 1.00 | [-0.01, 0.37] | [-0.04, 0.4] |
| beta_ land surface temperature | 0.0949 | 74.9 | 3,798 | 1.00 | [-0.14, 0.33] | [-0.18, 0.37] |
| beta_ landscape diversity | 0.00572 | 52.0 | 4,435 | 1.00 | [-0.18, 0.19] | [-0.22, 0.22] |
| beta_ edge density | −0.126 | 90.6 | 4,627 | 1.00 | [-0.28, 0.03] | [-0.31, 0.06] |

### **Supplementary Table 21.** Summary of Bayesian model evaluating the influence of urban environmental factors on bee nest demographic status.

| **Parameter** | **Median** | **Probability of Direction (%)** | **Effective Sample Size (ESS)** | **Rhat** | **90% CI** | **95% CI** |
| --- | --- | --- | --- | --- | --- | --- |
| alpha | 0.463 | 100.0 | 3,975 | 1.00 | [0.28, 0.64] | [0.24, 0.68] |
| beta_green cover | -0.0351 | 70.4 | 3,567 | 1.00 | [-0.15, 0.07] | [-0.17, 0.10] |
| beta_ land surface temperature | -0.122 | 95.9 | 3,707 | 1.00 | [-0.23, -0.01] | [-0.25, 0.01] |
| beta_ landscape diversity | 0.0842 | 89.9 | 3,281 | 1.00 | [-0.03, 0.19] | [-0.05, 0.22] |
| beta_ edge density | -0.0802 | 89.7 | 3,946 | 1.00 | [-0.18, 0.02] | [-0.20, 0.04] |

### **Supplementary Table 22.** Summary of Bayesian model evaluating the influence of urban environmental factors on wasp nest demographic status.

| **Parameter** | **Median** | **Probability of Direction (%)** | **Effective Sample Size (ESS)** | **Rhat** | **90% CI** | **95% CI** |
| --- | --- | --- | --- | --- | --- | --- |
| alpha | 0.313 | 96.8 | 4,070 | 1.00 | [0.04, 0.61] | [-0.02, 0.68] |
| beta_green cover | 0.0315 | 63.8 | 4,174 | 1.001 | [-0.11, 0.19] | [-0.14, 0.22] |
| beta_ land surface temperature | -0.0312 | 60.1 | 2,839 | 1.00 | [-0.23, 0.18] | [-0.26, 0.22] |
| beta_ landscape diversity | -0.0237 | 59.9 | 5,562 | 1.00 | [-0.19, 0.13] | [-0.22, 0.16] |
| beta_ edge density | 0.0415 | 67.7 | 6,580 | 0.999 | [-0.10, 0.19] | [-0.13, 0.22] |

### **Supplementary Table 23.** Summary of Bayesian model assessing the effect of urbanisation on bee community beta diversity.

| Parameter | Median | Probability of Direction (%) | Effective Sample Size (ESS) | Rhat | 90% CI | 95% CI |
| --- | --- | --- | --- | --- | --- | --- |
| alpha | 0.513 | 100 | 400 | 1.01 | [0.43, 0.59] | [0.38, 0.63] |
| beta_impervious surface cover | 0.0386 | 100 | 3,131 | 0.999 | [0.03, 0.05] | [0.02, 0.05] |
| beta_geographic distance | 0.0296 | 100 | 3,229 | 1.00 | [0.02, 0.04] | [0.02, 0.04] |

### **Supplementary Table 24.** Summary of Bayesian model assessing the effect of urbanisation on bee community turnover.

| Parameter | Median | Probability of Direction (%) | Effective Sample Size (ESS) | Rhat | 90% CI | 95% CI |
| --- | --- | --- | --- | --- | --- | --- |
| alpha | 0.454 | 100 | 827 | 1.01 | [0.36, 0.53] | [0.33, 0.56] |
| beta_impervious surface cover | 0.0230 | 99.6 | 4,218 | 1.00 | [0.01, 0.04] | [0.01, 0.04] |
| beta_geographic distance | −0.0353 | 100 | 3,660 | 1.00 | [-0.05, -0.02] | [-0.05, -0.02] |

### **Supplementary Table 25.** Summary of Bayesian model assessing the effect of urbanisation on bee community nestedness.

| Parameter | Median | Probability of Direction (%) | Effective Sample Size (ESS) | Rhat | 90% CI | 95% CI |
| --- | --- | --- | --- | --- | --- | --- |
| alpha | −0.210 | 99.7 | 912 | 1.00 | [-0.28, -0.14] | [-0.3, -0.12] |
| beta_impervious surface cover | −0.00232 | 59.1 | 2,904 | 1.00 | [-0.02, 0.02] | [-0.02, 0.02] |
| beta_geographic distance | 0.0432 | 100 | 2,826 | 1.00 | [0.02, 0.06] | [0.02, 0.07] |

### **Supplementary Table 26.** Summary of Bayesian model assessing the effect of urbanisation on wasp community beta diversity.

| Parameter | Median | Probability of Direction (%) | Effective Sample Size (ESS) | Rhat | 90% CI | 95% CI |
| --- | --- | --- | --- | --- | --- | --- |
| alpha | 0.408 | 99.6 | 1,237 | 1.00 | [0.23, 0.57] | [0.17, 0.61] |
| beta_impervious surface cover | −0.00879 | 75.4 | 3,974 | 1.00 | [-0.03, 0.01] | [-0.03, 0.02] |
| beta_geographic distance | 0.0113 | 82.2 | 3,242 | 1.00 | [-0.01, 0.03] | [-0.01, 0.03] |

### **Supplementary Table 27.** Summary of Bayesian model assessing the effect of urbanisation on wasp community turnover.

| Parameter | Median | Probability of Direction (%) | Effective Sample Size (ESS) | Rhat | 90% CI | 95% CI |
| --- | --- | --- | --- | --- | --- | --- |
| alpha | 0.706 | 100 | 828 | 1.00 | [0.57, 0.82] | [0.52, 0.84] |
| beta_impervious surface cover | −0.00205 | 56.9 | 4,044 | 1.00 | [-0.02, 0.02] | [-0.03, 0.02] |
| beta_geographic distance | −0.0206 | 97.6 | 3,173 | 1.00 | [-0.04, 0] | [-0.04, 0] |

### **Supplementary Table 28.** Summary of Bayesian model assessing the effect of urbanisation on wasp community nestedness.

| Parameter | Median | Probability of Direction (%) | Effective Sample Size (ESS) | Rhat | 90% CI | 95% CI |
| --- | --- | --- | --- | --- | --- | --- |
| alpha | −0.111 | 85.5 | 1,313 | 1.00 | [-0.28, 0.07] | [-0.32, 0.11] |
| beta_impervious surface cover | 0.0161 | 77.4 | 3,634 | 1.00 | [-0.02, 0.05] | [-0.02, 0.06] |
| beta_geographic distance | 0.0567 | 99.7 | 3,648 | 0.999 | [0.02, 0.09] | [0.02, 0.1] |

### **Supplementary Table 29.** Summary of Bayesian model evaluating the influence of urban environmental factors on bee community beta diversity.

| Parameter | Median | Probability of Direction (%) | Effective Sample Size (ESS) | Rhat | 95% CI | 90% CI |
| --- | --- | --- | --- | --- | --- | --- |
| alpha | 0.514 | 100 | 752 | 1.00 | [0.43, 0.59] | [0.45, 0.56] |
| beta_landscape diversity | −0.0200 | 99.9 | 3,799 | 1.00 | [-0.03, -0.01] | [-0.03, -0.01] |
| beta_land surface temperature | 0.0173 | 99.5 | 3,264 | 0.999 | [0, 0.03] | [0.01, 0.03] |
| beta_edge density | −0.00414 | 74.1 | 3,892 | 1.00 | [-0.02, 0.01] | [-0.01, 0.01] |
| beta_green cover | −0.0341 | 100 | 3,179 | 1.00 | [-0.05, -0.02] | [-0.05, -0.02] |
| beta_geographic distance | 0.0348 | 100 | 3,128 | 1.00 | [0.02, 0.05] | [0.02, 0.05] |

### **Supplementary Table 30.** Summary of Bayesian model evaluating the influence of urban environmental factors on bee community turnover.

| Parameter | Median | Probability of Direction (%) | Effective Sample Size (ESS) | Rhat | 95% CI | 90% CI |
| --- | --- | --- | --- | --- | --- | --- |
| alpha | 0.455 | 100 | 626 | 1.01 | [0.35, 0.57] | [0.38, 0.54] |
| beta_landscape diversity | −0.00623 | 77.0 | 4,876 | 1.00 | [-0.02, 0.01] | [-0.02, 0.01] |
| beta_land surface temperature | 0.0272 | 100 | 4,551 | 1.00 | [0.01, 0.04] | [0.01, 0.04] |
| beta_edge density | −0.00606 | 77.1 | 4,693 | 0.999 | [-0.02, 0.01] | [-0.02, 0.01] |
| beta_green cover | −0.0313 | 100 | 4,828 | 1.00 | [-0.05, -0.01] | [-0.05, -0.02] |
| beta_geographic distance | −0.0375 | 100 | 4,252 | 1.00 | [-0.06, -0.02] | [-0.05, -0.02] |

### **Supplementary Table 31.** Summary of Bayesian model evaluating the influence of urban environmental factors on bee community nestedness.

| Parameter | Median | Probability of Direction (%) | Effective Sample Size (ESS) | Rhat | 95% CI | 90% CI |
| --- | --- | --- | --- | --- | --- | --- |
| alpha | −0.210 | 99.8 | 732 | 1.01 | [-0.3, -0.1] | [-0.27, -0.14] |
| beta_landscape diversity | 0.0120 | 87.5 | 4,562 | 1.00 | [-0.01, 0.03] | [-0.01, 0.03] |
| beta_land surface temperature | −0.0352 | 99.9 | 4,085 | 1.00 | [-0.05, -0.02] | [-0.05, -0.02] |
| beta_edge density | −0.0121 | 89.2 | 3,401 | 1.00 | [-0.03, 0.01] | [-0.03, 0] |
| beta_green cover | 0.0285 | 99.6 | 3,122 | 1.00 | [0.01, 0.05] | [0.01, 0.05] |
| beta_geographic distance | 0.0460 | 100 | 3,713 | 1.00 | [0.02, 0.07] | [0.03, 0.06] |

### **Supplementary Table 32.** Summary of Bayesian model evaluating the influence of urban environmental factors on wasp community beta diversity.

| Parameter | Median | Probability of Direction (%) | Effective Sample Size (ESS) | Rhat | 95% CI | 90% CI |
| --- | --- | --- | --- | --- | --- | --- |
| alpha | 0.420 | 100 | 1,070 | 1.00 | [0.23, 0.61] | [0.26, 0.57] |
| beta_landscape diversity | 0.0383 | 99.8 | 3,305 | 1.00 | [0.01, 0.06] | [0.02, 0.06] |
| beta_land surface temperature | 0.0394 | 99.9 | 3,888 | 1.00 | [0.01, 0.06] | [0.02, 0.06] |
| beta_edge density | −0.0132 | 87.9 | 4,175 | 1.00 | [-0.03, 0.01] | [-0.03, 0.01] |
| beta_green cover | −0.0265 | 97.8 | 4,268 | 1.00 | [-0.05, 0] | [-0.05, 0] |
| beta_geographic distance | 0.0130 | 86.1 | 4,000 | 1.00 | [-0.01, 0.04] | [-0.01, 0.03] |

### **Supplementary Table 33.** Summary of Bayesian model evaluating the influence of urban environmental factors on wasp community turnover.

| Parameter | Median | Probability of Direction (%) | Effective Sample Size (ESS) | Rhat | 95% CI | 90% CI |
| --- | --- | --- | --- | --- | --- | --- |
| alpha | 0.711 | 100 | 956 | 1.00 | [0.52, 0.86] | [0.57, 0.83] |
| beta_landscape diversity | 0.0188 | 92.0 | 4,327 | 1.00 | [-0.01, 0.05] | [0, 0.04] |
| beta_land surface temperature | 0.0237 | 97.2 | 3,925 | 1.00 | [0, 0.05] | [0, 0.04] |
| beta_edge density | −0.00512 | 67.3 | 4,607 | 1.00 | [-0.03, 0.02] | [-0.02, 0.01] |
| beta_green cover | −0.0204 | 93.5 | 4,289 | 1.00 | [-0.05, 0.01] | [-0.04, 0] |
| beta_geographic distance | −0.0190 | 97.0 | 4,237 | 1.00 | [-0.04, 0] | [-0.04, 0] |

### **Supplementary Table 34.** Summary of Bayesian model evaluating the influence of urban environmental factors on wasp community nestedness.

| Parameter | Median | Probability of Direction (%) | Effective Sample Size (ESS) | Rhat | 95% CI | 90% CI |
| --- | --- | --- | --- | --- | --- | --- |
| alpha | −0.117 | 86.8 | 1,539 | 1.00 | [-0.35, 0.1] | [-0.3, 0.07] |
| beta_landscape diversity | 0.00376 | 57.0 | 5,213 | 1.00 | [-0.04, 0.05] | [-0.03, 0.04] |
| beta_land surface temperature | −0.0689 | 99.9 | 5,191 | 1.00 | [-0.11, -0.03] | [-0.1, -0.03] |
| beta_edge density | −0.0328 | 95.3 | 5,160 | 1.00 | [-0.07, 0.01] | [-0.06, 0] |
| beta_green cover | 0.0214 | 83.5 | 5,502 | 0.999 | [-0.02, 0.06] | [-0.01, 0.06] |
| beta_geographic distance | 0.0610 | 99.8 | 4,891 | 1.00 | [0.02, 0.1] | [0.03, 0.1] |

### **Supplementary Methods 1.** Detailed protocol for species identification using COI barcoding.

DNA was extracted from emerged individuals for species identification using mitochondrial cytochrome oxidase I (COI) barcoding. A ~650 bp region of the mitochondrial cytochrome-c oxidase subunit-I gene was amplified with the primers LCO-1490 and HCO-2198 (Folmer et al., 1994). PCR reactions were carried out in 10 µL volumes consisting of 1 x PCR buffer containing 1.5 mM MgCl_2_ (Promega, Madison, WI, USA), 200 µM of each dNTP, 0.4 µM of each primer, 1.5 U Taq-polymerase (Promega) and 2 µL of template DNA (ca. 25-50 ng). PCRs were performed with a thermocycler (Biometra, Göttingen, Germany) under the following thermal regime: 3 min at 94°C, followed by 35 cycles of 30 s at 94°C, 45 s of annealing at 50°C and 1 min at 72°C for elongation and a final elongation step at 72°C for 8 min. PCR products were purified using an ExoSAP-IT PCR product cleanup kit (Affymetrix, Santa Clara, CA, USA) and Sanger sequenced using the LCO-1490 primer. Sequences were trimmed and quality-checked in Geneious v. 7.1.9 (https://www.geneious.com) and species were identified using BLAST searches against NCBI GenBank and BOLD, with a ≥98% identity threshold (Ratnasingham & Hebert, 2007).

### **Supplementary Methods 2.** Partitioning beta diversity into turnover and nestedness components.

According to the framework proposed by Baselga (2010, 2012), Jaccard beta diversity (β_jac_) for a pair of sites can be partitioned into turnover (β_jtu_, species replacement between sites) and nestedness composition (β_jne_, species losses or gains between sites) with the following equations:

$$\beta_{\mathrm{jac}}=\frac{b+c}{a+b+c}$$

( 1 )

$$\beta_{jtu}=\frac{2\times\min\left( b,c \right)}{a+2\times\min\left( b,c \right)}$$

( 2 )

$$\beta_{jne}=\beta_{jac}-\beta_{jtu}=\frac{\left( b+c \right)}{\left( a+b+c \right)}-\frac{2\times\min\left( b,c \right)}{a+2\times\min\left( b,c \right)}$$

( 3 )

where *a* is the number of species present in both sites, *b* is the number of species present in the first site but not in the second, and *c* is the number of species present in the second site, but not in the first.

### **Supplementary Methods 3.** Calculating Shannon index of landscape diversity and edge density.

Landscape diversity was calculated using the Shannon-Wiener index (H'):

$$H^{'}=-\sum_{i=1}^{S} p_{i} \ln\left( p_{i} \right)$$

( 4 )

where *pᵢ* represents the proportion of each land cover type *i* and s represents the number of land cover classes.

Landscape fragmentation was calculated as:

$$LF=\frac{E}{A}$$

( 5 )

where E represents the total length of patch edges and A represents the total area.
